# Specific Bacterial Taxa and Their Metabolite, DHPS, Linked to Alzheimer’s Disease, Parkinson’s Disease, and Amyotrophic Lateral Sclerosis

**DOI:** 10.1101/2024.10.13.615818

**Authors:** Courtney Jayde Christopher, Katherine Hope Morgan, Christopher Mahone Tolleson, Randall Trudell, Roberto Fernandez-Romero, Lexis Rice, Blessing A Abiodun, Zane Vickery, Katarina Jones, Brittni Morgan Woodall, Christopher Nagy, Piotr Andrzej Mieczkowski, Shawn R Campagna, Gregory Bowen, Joseph Christopher Ellis

**Affiliations:** Department of Chemistry, University of Tennessee, 1420 Circle Drive, Knoxville, TN 37996; College of Nursing, University of Tennessee, 1412 Circle Drive, Knoxville, TN 37996; University of Tennessee Medical Center, Division of Neurology, 1924 Alcoa Hwy, Knoxville, TN 37920; The Pat Summitt Clinic at The University of Tennessee Medical Center, 1932 Alcoa Hwy., Suite C-150, Knoxville, TN 37920; Department of Biochemistry & Cellular and Molecular Biology, University of Tennessee, 1311 Cumberland Ave., Knoxville, TN 37916; Department of Chemistry, University of Tennessee, Department of Chemistry, University of Tennessee, 522 Buehler Hall, 1420 Circle Drive, Knoxville, TN 37996; High Throughput Sequencing Facility; University of North Carolina at Chapel Hill, School of Medicine, 1153 Genome Science Building, SOM CB#7102, Chapel Hill, NC 27599; Department of Genetics, University of North Carolina at Chapel Hill, School of Medicine, 4256 Genome Science Building, SOM CB#7264, Chapel Hill, NC 27599; Department of Chemistry and Biological and Small Molecule Mass Spectrometry Core, University of Tennessee, 611 Buehler Hall, 1420 Circle Drive, Knoxville, TN 37996-1600; Integrated Genomics Cores; University of North Carolina at Chapel Hill, 4252 Genome Sciences Building, 250 Bell Tower Dr., Chapel Hill, NC 27514; NetEllis, and Department of Medicine at the UT Graduate School of Medicine, Knoxville, TN 37920

## Abstract

Neurodegenerative diseases (NDDs) are multifactorial disorders frequently associated with gut dysbiosis, oxidative stress, and inflammation; however, the pathophysiological mechanisms remain poorly understood. We investigated bacterial and metabolic dyshomeostasis in the gut microbiome associated with early disease stages across three NDDs, amyotrophic lateral sclerosis (ALS), Alzheimer’s Disease (AD), Parkinson’s Disease (PD), and healthy controls (HC) and discovered a previously unrecognized link between a microbial-derived metabolite with an unknown role in human physiology, 2,3-dihydroxypropane-1-sulfonate (DHPS), and NDDs. DHPS was downregulated in AD, ALS, and PD, while *Eubacterium* and *Desulfovibrio,* capable of metabolizing this metabolite,^1–4^ were increased in all disease cohorts. Additionally, select taxa within the Clostridia class had strong negative correlations to DHPS suggesting a potential role in DHPS metabolism. Hydrogen sulfide is a catabolic product of DHPS,^1,5^ and hydrogen sulfide promotes inflammation,^6–8^ oxidative stress,^9^ mitochondrial damage,^10^ and gut dysbiosis,^2,11^ known hallmarks of NDD. These findings suggest that cryptic sulfur metabolism via DHPS is a missing link in our current understanding of NDD onset and progression. To the best of our knowledge, we are the first to provide evidence of a conserved gut-brain axis linkage of specific bacterial taxa and their metabolism of DHPS shared by three neurodegenerative diseases.

## Main

The social and economic impact of neurodegenerative diseases (NDDs) such as Alzheimer’s (AD), amyotrophic lateral sclerosis (ALS), and Parkinson’s (PD) have devasting impacts on patients and families alike. In aggregate, neurological disorders cause disability, reduce lifespan, and are the second leading cause of death worldwide.^12^ A growing number of researchers are exploring the gut-brain axis in Alzheimer’s disease using the mouse model.^13^ However, the cause of neurodegenerative disease, broadly, and amyotrophic lateral sclerosis, Alzheimer’s Disease, Parkinson’s Disease, specifically, remain poorly understood.

The gut is in constant system-wide communication with organs throughout the body,^14^ and dyshomeostasis within the metabolome can contribute to intestinal and systemic inflammation, oxidative stress, and mitochondrial dysfunction which are involved in the pathology of NDDs.^15^ For example, hydrogen sulfide (H_2_S) is a key metabolite involved in sulfur metabolism with a myriad of beneficial physiological functions and is primarily produced by members of the *Desulfovibrio* genus.^2,16^ However, excess H_2_S promotes inflammation,^6–8^ oxidative stress,^9^ mitochondrial damage,^10^ and gut dysbiosis.^2,11^ Dyshomeostasis in sulfur metabolism has been well-documented in PD,^17–19^ and the host-microbe regulation of oxidized sulfur compounds in the metabolome is gaining recognition in health and disease.^20^

In this study, we collected fecal samples and investigated the human gut bacterial microbiome and metabolome of patients from three NDDs and a healthy controls (HC) cohorts using a multi-omics approach. Sampling occurred early in disease progression, at the first or second clinical appointment with a neurologist. These results demonstrate a previously unrecognized association between microbial-derived metabolite DHPS and NDDs. While the role of DHPS in human health is unknown, we found DHPS to be significantly downregulated in the stool metabolome of AD, ALS, and PD cohorts compared to controls. In addition, we observed that specific taxa responsible for DHPS production (*Eubacterium*) and catabolism (*Desulfovibrio)* were significantly increased in NDDs. For the first time, we have provided data directly linking DHPS to NDDs. These findings may suggest that DHPS is a missing link in the current understanding of sulfur dyshomeostasis in the gut associated with NDDs.

### Subjects Were in Early Diagnostic Stages

From summer 2021 to spring 2022, 48 participants enrolled, and 45 completed the study. Of these, 39 submitted stool samples (AD=5, ALS=11, PD=13, and HC=10). Participants were non-Hispanic Caucasians living within the 21-counties served by the University of Tennessee Medical Center (See Supplementary Table 1 for additional demographics) in Knoxville, Tennessee.

### Global Metabolomics

Untargeted metabolomics was used to investigate functional alterations in the gut metabolome associated with NDDs. We identified 164 metabolites based on exact mass and retention time and evaluated metabolic profile differences between each NDD and HC using PLS-DA, which revealed that each NDD had a unique gut metabolic profile compared to HC (Fig. 1). To determine metabolites altered by NDD, metabolites from each pairwise comparison with a fold change > |1.5| (p < 0.1) or VIP score > 1 were used for further analysis (Supplementary Table 2). These metabolites were also used to identify unique metabolic signatures for each NDD. This revealed 14, 20, and 9 unique markers for AD, ALS, and PD, respectively. Moreover, we identified 19 metabolic markers, which were consistently altered across all three NDDs in this study (Supplementary Table 3).

**Figure 1.**
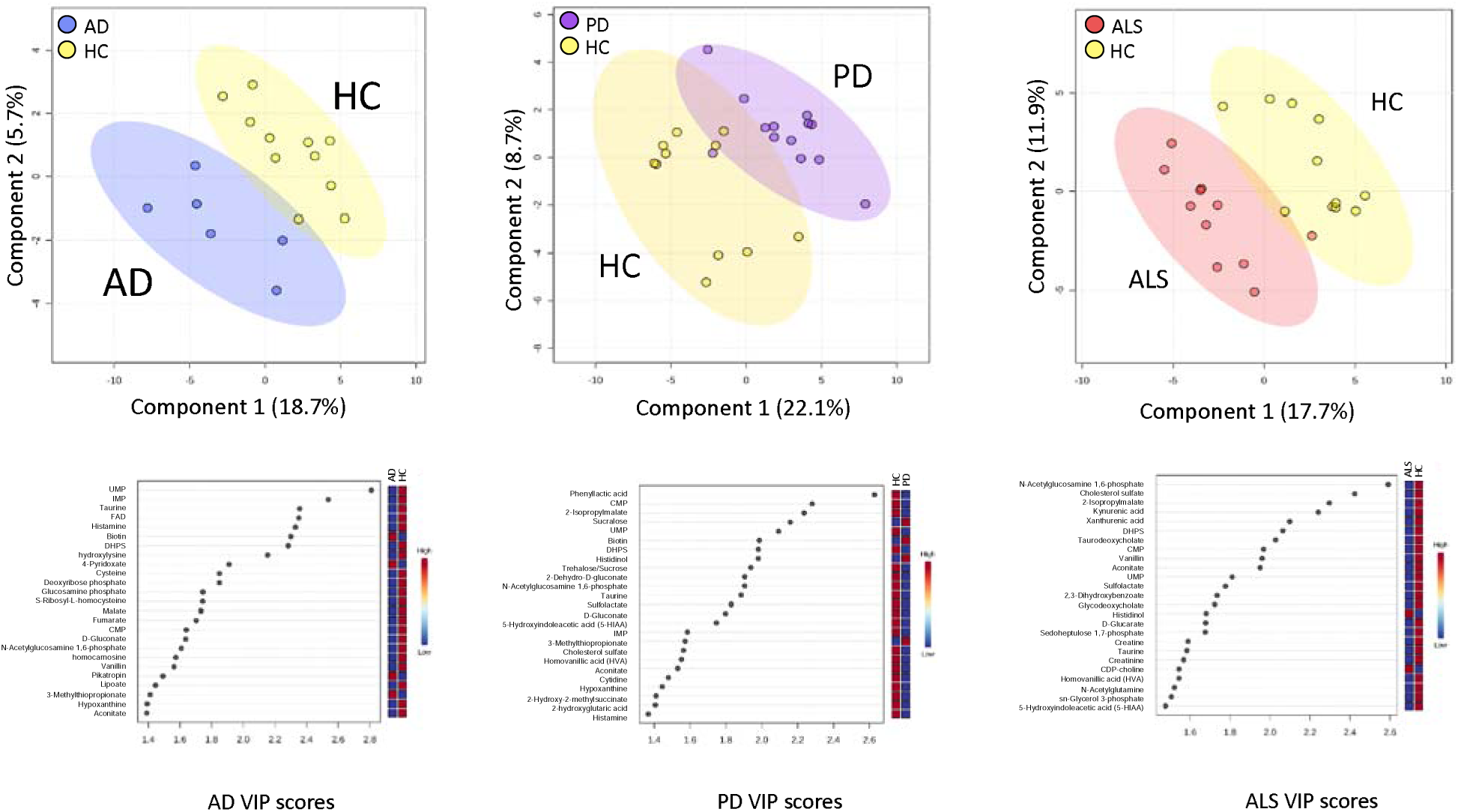
Partial least squares discriminant analysis (PLS-DA) comparing metabolomes associated with NDD and HC. Individual participants are shown with respective confidence intervals; HC samples are yellow; AD samples are blue; PD samples are purple; ALS samples are shown in red. D-F) Variable importance in projection (VIP) scores for each corresponding PLS-DA model top show the top 25 metabolites contributing most to the differences in metabolic profiles.

#### AD

Based on the PLS-DA model between the AD and HC metabolomes, the overall metabolic profile of the gut in AD patients was distinctly different from HC as seen by the clustering and separation of groups (Fig. 1). Metabolites with the highest VIP scores, thereby contributing most to global metabolome differences, were UMP (2.8), IMP (2.5), and taurine (2.4). We observed that 29% of identified metabolites in the gut were altered by AD, and 14 of these metabolites were unique to AD. Notably, the primary catabolic product of vitamin B_6_ (4-pyridoxate), S-ribosyl-L-homocysteine, 3-methylphenylacetic acid, and metabolites involved in fatty acid and energy metabolism (carnitine, hydroxylysine, FMN) were unique to AD. Metabolic profile differences in AD demonstrated altered sulfur metabolism, with downregulation of cysteine, taurine, methionine, homocysteine, and DHPS, while sulfur containing B vitamins, biotin and pyridoxine, were increased in the AD cohort compared to HC (Fig. 2).

**Figure 2.**
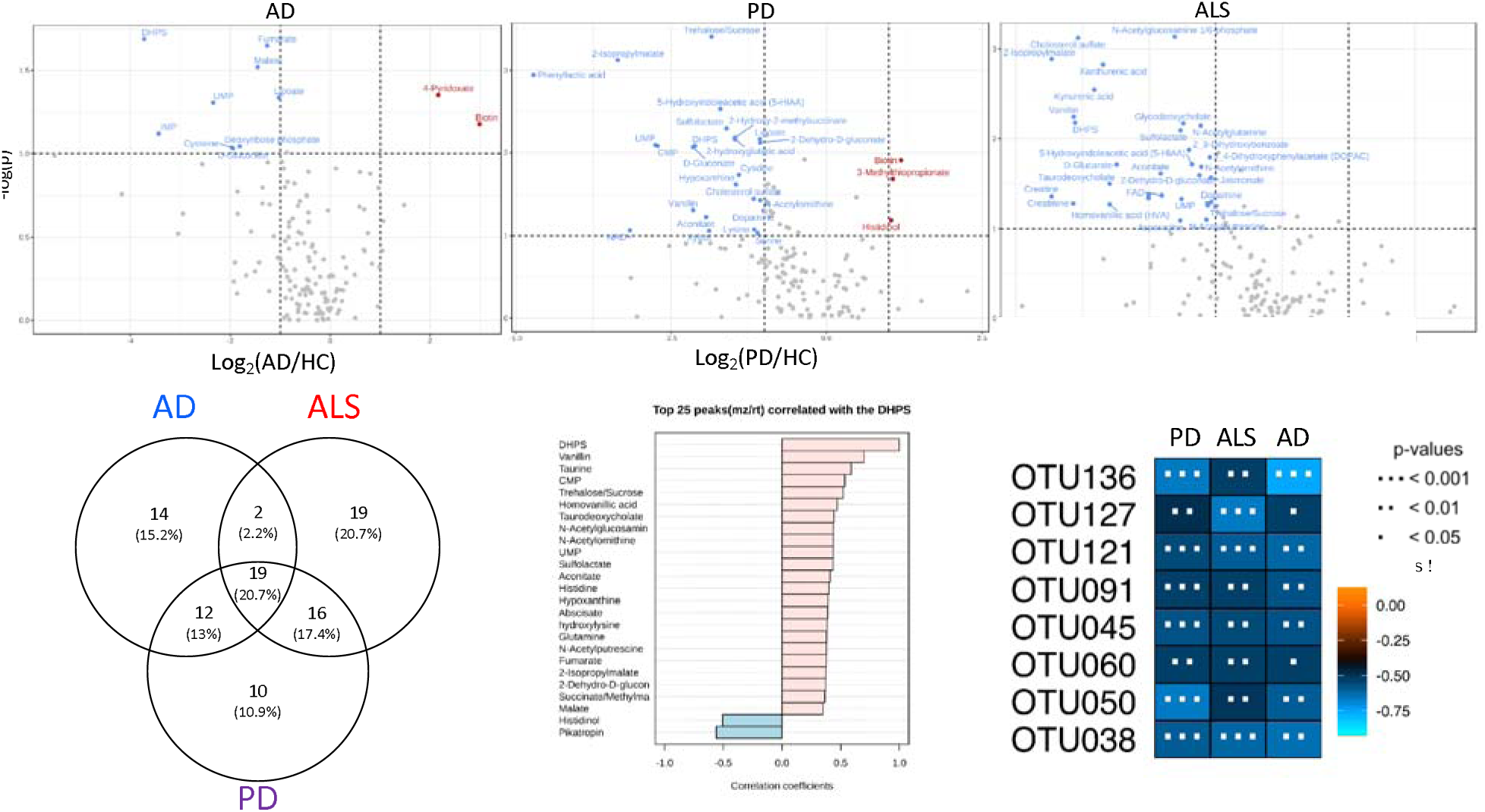
There were 19 metabolic markers for all NDD in this study, as well as unique markers for each disease. DHPS was one of the 19 markers for NDD as was significantly decreased in NDD cohorts. DHPS was strongly correlated to acetylated amino acids and taurine. A-C) volcano plots showing metabolites significantly different between cohorts. The x-axis is the log_2_ foldchange and y-axis is the −log p value. The following cutoffs were used: fold change > |1.5| (p < 0.1). D). Venn diagram showing unique and shared metabolic markers. These metabolites had fold change > |1.5| (p < 0.1) or VIP score >1. E). Metabolites strongly correlated with DHPS in PD. F). Eight OTUs were significantly correlated with DHPS across all three NDDs. OTU136 (d_Bacteria;p_Firmicutes;c_Clostridia_;_;_;) OUT127 (d_Bacteria;p_Firmicutes;c_Clostridia;o_Clostridia;f_Hungateiclostridiaceae;_) OTU121 (d_Bacteria;p_Firmicutes;c_Clostridia;o_Oscillospirales;f_Oscillospiraceae;g_Oscillospir_a) OTU091 (d_Bacteria;p_Firmicutes;c_Clostridia;o_Oscillospirales;f_UCG-010;g_UCG-010) OTU045 (d_Bacteria;p_Firmicutes;c_Clostridia;o_Clostridia_vadinBB60_group;f_Clostridia_vadin_BB60_group;g_Clostridia_vadinBB60_group) OTU060 (d_Bacteria;p_Bacteroidota;c_Bacteroidia;o_Bacteroidales;f_Marinifilaceae;g_Odoribacter) OTU050 (d_Bacteria;p_Firmicutes;c_Clostridia;o_Oscillospirales;f_Oscillospiraceae;_) OTU038 (d_Bacteria;p_Firmicutes;c_Clostridia;o_Oscillospirales;f_[Eubacterium]_coprostanoligenes_group;g_[Eubacterium]_coprostanoligenes_group)

#### ALS

Similarly, the ALS cohort also exhibited global gut metabolome differences compared to HC, as shown in Figure 1. Metabolites contributing most to differences between cohorts were N-acetylglucosamine phosphate (VIP = 2.6), cholesterol sulfate (VIP = 2.4), 2-isopropylmalate (VIP = 2.3), and kynurenic acid (VIP = 2.2) (Fig. 2). Overall, 34% of identified metabolites were significantly altered in the ALS cohort: of these, histadinol and CDP-choline were increased, while all other metabolites were decreased. Similar to AD, we observed that ALS caused alterations in tryptophan metabolism, sulfur metabolism, and acetylated amino acids in the gut (Fig. 2). There were 19 metabolites unique to ALS, including metabolites involved in tryptophan metabolism (xanthurenic acid, kynurenic acid), vitamin B_6_ (pyridoxine), and energy, fatty acid, and amino acid metabolism (creatine, creatinine, arginine, CDP-choline, and glycodeoxycholate). Interestingly, creatine metabolism was unique to the ALS cohort, with no notable alterations in AD or PD cohorts.

#### PD

The most dramatic alterations in the gut metabolome were found when comparing the PD to HC cohort with 35% of identified metabolites being altered by PD (Fig. 1). Phenyllactic acid (2.6), CMP (2.3), 2-isopropylmalate (2.3), sucralose (2.2), and UMP (2.1) had the highest VIP scores contributing most to the global metabolome differences between the PD and HC cohorts in the PLS-DA model. Similar to both AD and ALS, PD caused alterations in sulfur, tryptophan, vitamin B, and acetylated amino acid metabolism. Other notable differences in the PD gut metabolome include downregulation of taurine, alanine, nicotinate and nicotinamide, and cysteine and methionine metabolism (Fig 2). Conversely, biotin, sucralose, 3-methylthiopropionate, and histidinol were upregulated in PD (Fig 2). There were 10 unique metabolic markers associated with PD including xylose, lysine, xylitol, and N-acetyl-beta-alanine. The most dramatic alterations in neurotransmitters (5-Hydroxyindoleacetic acid (5-HIAA), dopamine, 3,4-dihydroxyphenylacetate (DOPAC)) were observed in the gut metabolome of PD patients.

### Neurodegenerative disease metabolic markers

There were 19 metabolites altered by the three NDDs in the study, serving as metabolic markers for AD, ALS, and PD (Fig. 2). Many of these metabolites were involved in sulfur and vitamin B metabolism (DHPS, taurine, 3-methylthiopropionate, cholesterol sulfate, sulfolactate, and biotin). DHPS is a microbial-derived metabolite, and was on average 9x lower (*p* < 0.05) in the NDD cohorts compared to the HC cohort (Fig. 2). We also observed that DHPS had strong correlations to acetylated amino acids and metabolites involved in neurotransmitter, bile acid, and vitamin B metabolism (Fig. 2).

### Sequencing (16s rRNA)

The microbial composition of stool samples from NDD and HC patients were characterized via 16S amplicon sequencing. Differences in alpha diversity (Fig. 3) were only observed when comparing the PD and HC cohorts, with PD having higher alpha diversity (p = 0.02). Analysis of OTUs revealed phylum level differences in the abundance of Firmicutes in PD (53%) compared to HC (67%) stool, while Firmicutes displayed minimal differences in ALS and AD cohorts (Fig. 3). Phylum Actinobacteria was unchanged in AD but was increased in PD and ALS compared to HC patients. Differences within the Firmicutes phylum in PD could broadly be attributed to genus *Eubacterium*. There were six taxa significantly altered across all NDDs compared to HCs: *Desulfovibrio, Eubacterium, Akkermansia, Paludicola, Sellimonas, and Enterococcus*. (Fig. 3) All of these taxa were more abundant in the NDD cohorts than HC, except for *Enterococcus* which was higher in HC.

**Figure 3.**
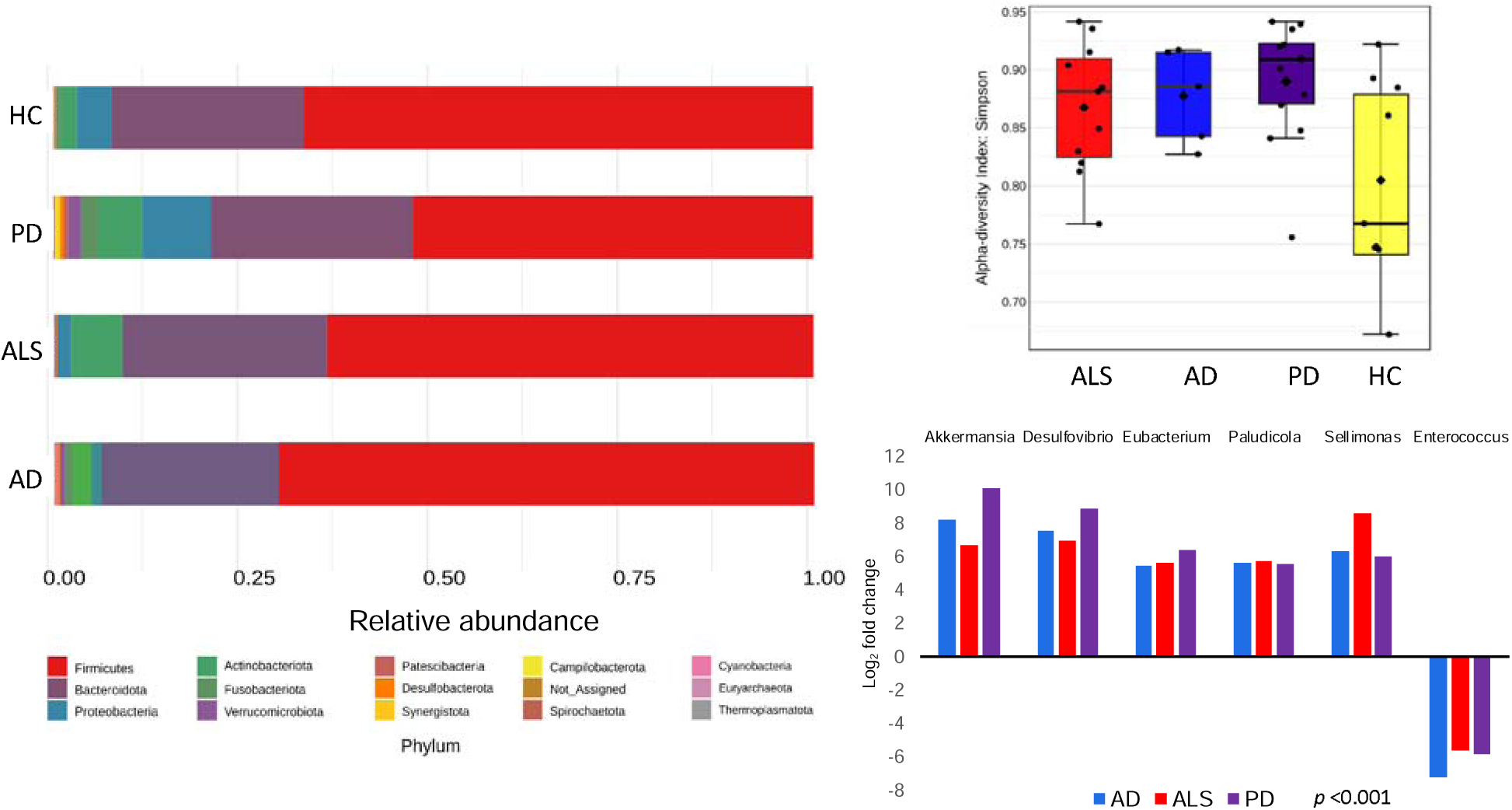
Gut microbial composition is altered in NDDs A) Phyla level differences in microbial composition, B) Alpha diversity was significantly greater in PD than HC cohort. There were no significant differences in the AD and ALS cohort compared to HC, C) log_2_ fold change of taxa significantly altered in across AD, ALS, and PD cohorts. Orange represents higher abundance in disease compared to HC while blue represents lower abundance in disease compared to HC.

To assess the relationship between gut microbial composition and function in NDDs, we identified OTUs correlated with metabolites that were significantly different in NDDs. Microbial-derived metabolite DHPS was consistently and significantly altered in NDDs and there were 33, 43, and 15 OTUs significantly correlated with DHPS in PD, AD, and ALS, respectively, all having negative correlations. Of these, eight OTUs were correlated with DHPS across PD, AD, and ALS, and all were part of the Firmicutes phylum (class *Clostridia*), except for one belonging to the Bacteroidetes phylum (Fig. 2).

## Discussion

### Cryptic sulfur metabolism in NDDs

All detected metabolites in the transsulfuration pathway were decreased in NDD cohorts, except for cystathionine. Rapid conversion of homocysteine to cystathionine could suggest increased fluxes in sulfur metabolism as previous work using longitudinal metabolome data and constraint-based modeling of gut microbial communities has shown increased flux in the transsulfuration pathway in PD.^18^ This pathway generates glutathione and H_2_S, which impact redox metabolism. Increased fluxes through sulfur metabolism may be indicative of excess H_2_S, which at low concentrations, is neuroprotective and cytoprotective. However, at high levels, H_2_S induces systemic inflammation,^6–8^ oxidative stress,^9^ mitochondrial damage,^10^ and gut dysbiosis.^2,11^ Our findings support the current hypothesis that the increased burden of oxidative stress causes mitochondrial dysfunction leading to neuroinflammation.

In addition to metabolites in the transsulfuration pathway, other sulfur-containing metabolites, DHPS and taurine, were significantly altered across NDDs. We found this to be intriguing as oxidized sulfur-containing small molecules represent an underappreciated class of unique, biologically relevant metabolites in host-microbiome interactions.^20^ It has been discovered recently that these molecules play a significant role in cellular signaling and impact disease-related phenotypes thereby bringing oxidized sulfur metabolites to the forefront of enhancing our understanding of the pathophysiological impact of host-microbiome interactions.^20–24^ Since the role of DHPS in NDDs is unknown, we identified metabolites and OTUs significantly correlated with DHPS. Five metabolites (taurine, vanillin, histidinol, *N*-acetylornithine, and sucrose/trehalose), and eight OTUs were significantly (p<0.01) correlated with DHPS in each NDD cohort. All of the significantly correlated OTUs belonged to phylum Firmicutes (orders Clostridia and Oscillospirales) except for one, which belonged to the phylum Bacteroidetes (genus *Odoribacter*). All of these microbes have been directly linked to metabolizing dietary sulfur to H_2_S.^4,25^ Together, the metabolites and OTUs correlated with DHPS suggest that DHPS is involved in cryptic sulfur host-microbiome metabolism in NDDs.

### DHPS microbial metabolism in NDDs

Frommeyer and colleagues annotated genes required for a novel bacterial transaldolase pathway by which select gut microbes (*Enterococcus*, *Clostridium*, *Desulfovibrio*, and *Eubacterium* strains) can ferment dietary sulfoquinovose (SQ) to H_2_S with DHPS as a transient intermediate.^2^ SQ is obtained through the intake leafy greens and vegetables containing the lipid sulfoquinovosyl diacylglycerol (SQDG), which is a highly abundant sulfur containing metabolite common in photosynthetic plants and cyanobacteria. In addition, Hanson and colleagues^2^ reported that DHPS contributes to intestinal H_2_S production and that *Eubacterium rectale (E. rectale)* is the primary DHPS producer, while *Desulfovibrio* and *Bilophila are* the primary DHPS consumers, in fecal microcosms from human vegetarians. They identified genes necessary for microbial catabolism of diet-acquired SQ to H_2_S and analyzed the expression of these genes across stool metatranscriptomes of 123 healthy, 28 ulcerative colitis, and 50 Crohn’s Disease patients to identify the overall impact on health.^2^ However, they found no significant differences in expression of these pathways between cohorts with bowel disease and healthy individuals.^2^

*Eubacterium rectale* plays a significant role in promoting inflammation and damaging tissue in the gut.^26^ We found that the genus *Eubacterium* was increased in PD, ALS, and AD, and it was one of the eight OTUs strongly correlated with DHPS across all three diseases (Fig. 3). Additionally, sulfate-reducing bacteria (SRB), such as *Desulfovibrio,* generate H_2_S, and have previously been implicated in the development and severity of PD.^17^ Here, we observed *Desulfovibrio* were significantly altered in PD, ALS, and AD and were more abundant in patients with NDDs (Fig. 3). Additionally, taurine was strongly correlated with DHPS and significantly decreased in NDDs and can also be obtained through dietary sources and degraded by *Desulfovibrio and Bilophilia* to acetyl-CoA and H_2_S. Other studies have also implicated *Bilophilia wadsworthia* in PD altering microbial sulfur metabolism, yet the etiology remains elusive.^18,27–29^ Disrupted sulfur metabolism in NDDs could result in increased flux through DHPS and taurine, promoting overgrowth of SRB and excess H_2_S. This may explain why DHPS producing and consuming microbes are increased in NDDs, while DHPS is decreased.

While we do not have dietary information from participants, it has become fairly accepted that the gut microbiome composition and function in NDDs are influenced by dietary habits and nutrition.^30^ Since diet-acquired SQDG from leafy greens can be bioconverted to DHPS by select microbes in the gut, we hypothesize that diet corresponds to DHPS and sulfur homeostasis in NDDs.

Previous studies have laid the foundation for investigating DHPS in human metabolism by elucidating the biochemical mechanism of DHPS production and degradation in the human gut and directly linking dietary sulfonates to H_2_S production;^2–4^ however, the data did not demonstrate an association between DHPS, or the genes involved in DHPS metabolism, and human health outcomes. Recently, Zaparte, Christopher and colleagues^31^ provided the first report of direct DHPS detection in human stool samples and highlighted a compelling connection between DHPS and gut metabolic dysregulation in people who use e-cigarette and/or combustible tobacco/marijuana smoke. They observed that DHPS was significantly higher (20x) in the control cohort compared to the smoking cohorts and noted strong correlations between DHPS and vitamin B, acetylated amino acids, bile acids, and cholesterol metabolism. This is consistent with the DHPS findings outlined herein, demonstrating a conserved trend across independent datasets that DHPS abundance in human stool is associated with overall health status, supporting the proposal of DHPS as a missing link in the pathophysiology of inflammation as it may be a key regulator of inflammation and overall human health.

### Conclusion

In this study, we investigated bacterial and metabolic dyshomeostasis in the gut associated with AD, ALS, and PD using an integrative cross-omics approach. Herein, we present the first linkage of DHPS, a microbial-derived metabolite, to NDDs, and suggest that it contributes to dyshomeostasis in sulfur metabolism, since DHPS consuming and degrading microbes are more abundant in NDDs while DHPS is decreased (Fig. 4). We hypothesize that the decrease in DHPS in NDDs may be a result of increased flux through the recently identified transaldolase pathway in which dietary sulfonates are metabolized to DHPS and H_2_S by select gut microbes, including *Eubacterium* and *Desulfovibrio*. Excess H_2_S, resulting from increased flux through DHPS, contributes to mitochondrial dysfunction, inflammation, oxidative stress, and gut dysbiosis in NDDs. These findings suggest that cryptic sulfur metabolism via DHPS is a missing link in our current understanding of NDD onset and progression.

**Figure 4.**
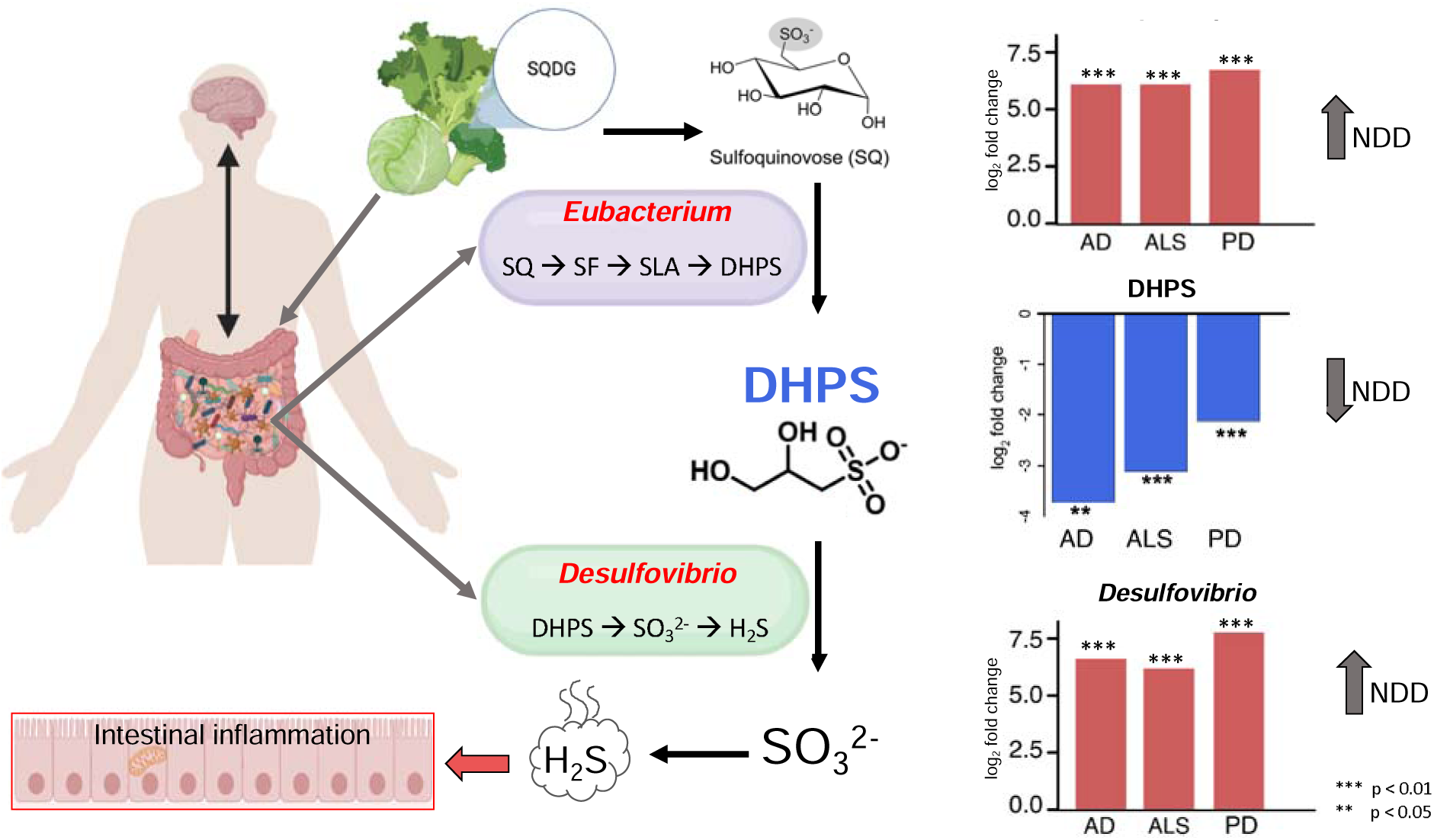
DHPS can be obtained through dietary consumption of leafy greens and degraded to H_2_S. The abundance of DHPS producing and consuming microbes are increased in NDD while DHPS is decreased in NDD suggesting the rapid degradation of DHPS to yield H_2_S. Other microbes (*Enterococcus* and *Clostridium*) can ferment dietary sulfoquinovose (SQ) to H_2_S with DHPS as a transient intermediate, as well. Figure was created using biorender.com and adapted from Hanson et al. ^2^

### Limitations

This study lays the foundation for future discoveries on the role of specific bacterial taxa and their metabolite DHPS however, there are some notable limitations. First, all patients were sampled in the same geographic location of Eastern Tennessee and the surrounding Appalachian Mountains. In addition, all the patients were Caucasian. Combined, this limits the generalizability of the conclusions. Finally, while this study describes a remarkably well conserved and novel association across different neurodegenerative diseases, this study is limited by not being able to define these findings as causative given the exploratory nature of untargeted metabolomics.

## Supporting information

Abstract Art

## Methods

### Study Population

This study was approved by the institutional review board of the University of Tennessee Health Science Center for the protection of human subjects. Participants were recruited during their first or second visit to a movement disorder specialist for ALS, AD, or PD from the 21-county regional service area of the University of Tennessee Medical Center, in Knoxville, Tennessee.

Participation was voluntary and only those who were willing and able to complete the study activities (with a caregiver if needed) were included. Participants were excluded if they had received antibiotics in the previous six months. Further inclusion criteria for each cohort were as follows: 1) For AD: 65 years of age or older, met all clinical criteria for mild-to-moderate AD according to National Institute on Aging-Alzheimer’s Association criteria,^1^ were otherwise in good health based on medical history and clinical examination; 2) ALS: age of 30 years or older at enrollment, and met the El Escorial Criteria^2^ for definite, probable, or possible ALS. 3) PD: age 50 to 85 years at enrollment, diagnosed within the last 5 years, absence of PD medications with the exception of MAO-B inhibitors, absence of dyskinesia or motor fluctuations, modified Hoehn and Yahr Stage^3^ <2.5 (symptoms ranging from unilateral involvement only [Stage 1] to mild bilateral involvement with recovery on a pull test [Stage 2.5]); and 4) HC: age 30 or older, were age matched (+/-4 years) and sex-matched, in good health based on medical history and clinical examination. HC age 65 years or older completed the Neuropsychological Battery from the National Alzheimer’s Coordinating Center^4^ and the Cognivue® computerized cognitive assessment prior to enrollment to demonstrate normal cognition.

### Sample Collection, Preservation, DNA Extraction

Samples were collected at home by participants within one week of enrollment. Participants were instructed to place stool from the same specimen into two vials as follows: 1) the DNA Genotek® gut microbiome DNA collection kit and 2) a 50-mL sterile tube, which was immediately frozen at home at −20 C. Stool samples were delivered frozen on ice by same-day courier service, to the laboratory, where they were aliquoted and stored at −80 C. Nucleic acid extraction was conducted in batches, using Zymo *Quick*-DNA Fecal/soil Microbe Kit, and extractions were frozen at −80 C, then shipped on dry ice to the University of North Carolina (UNC) Chapel Hill High Throughput Sequencing Facility (HTSF) for 16s rRNA/DNA sequencing.

### Metabolomics

#### Metabolite extractions

All samples were extracted and analyzed at the Biological and Small Molecule Mass Spectrometry Core at the University of Tennessee Knoxville (RRID: SCR_021368). Stool samples were kept at 4(, pre-weighed and homogenized. The exact mass of stool used for sample was recorded and used for normalization. Roughly 50 mg of stool were aliquoted and water-soluble metabolites were extracted using an acidic acetonitrile extraction procedure adapted from Rabinowitz and Kimball^5^ using 1.5 mL of 4:4:2 acetonitrile: methanol: water with 0.1 M formic acid. ^6,7^ All solvents were HPLC grade. The supernatant collected from each sample were dried under nitrogen, then 300 µL of LC-MS grade water was added to each sample before mass spectral analysis.

#### UHPLC-HRMS

A previously described ultra high-performance liquid chromatography high resolution mass spectrometry (UHPLC-HRMS) method was used for untargeted metabolomics analysis^8^ using an UltiMate 3000 RS autosampler (Dionex, Sunnyvale, CA, USA), Synergi 2.6 µm Hydro RP column (100 mm × 2.1 mm, 100 Å; Phenomenex, Torrance, CA, USA), an UltiMate 3000 pump (Dionex), and Exactive Plus Orbitrap mass spectrometer (Thermo Fisher Scientific, Waltham, MA, United States). A previously described 25 min gradient elution, reverse phase ion-paring method with a water:methanol solvent system, and a tributylamine ion pairing reagent was used for chromatographic separation.^9^ Metabolites were ionized via negative mode electrospray ionization (ESI) prior to full scan mass spectral analysis as previously described.^8^

#### Metabolomics data processing

Raw mass spectral files were converted to mzML files using a package from ProteoWizard, msConverter. ^10^ All mzML files were imported into an open-source software, metabolomics analysis and visualization engine (El-MAVEN) where metabolites were manually identified using an in-house library based on exact mass (±5 ppm) and retention time (±2 min). ^6,11,12^ Metabolite peaks were integrated and raw peak intensities for identified metabolites were exported from El-MAVEN to a csv file. Prior to statistical analysis, raw spectral data were normalized by mass of sample used for extraction.

#### Statistical Analysis

The normalized data were imported into MetaboAnalyst 5.0 and were filtered via interquartile range (IQR), log transformed, and Pareto scaled ^13–15^ Partial least squares discriminant analysis (PLS-DA) were performed in MetaboAnalyst 5.0 where variable importance in projection (VIP) scores were assigned to each metabolite to indicate the importance of each metabolite in contributing to the separation between experimental groups. VIP scores > 1 indicate that a metabolite significantly contributes to the separation of groups in the PLS-DA model. Venn diagrams were constructed using metabolites with VIP scores > 1 from pairwise group comparisons. MetaboAnalyst 5.0 was also used to generate volcano plots to visualize metabolites significantly different between groups.

#### 16s rRNA sequencing

In this study, we implemented a detailed two-step protocol for 16S rRNA gene amplification,^16^ prioritizing high precision and reproducibility across numerous samples. The initial PCR reaction was meticulously configured, utilizing specific volumes of Kapa Enhancer, Kapa Buffer A (KAPA Robust 2G-Roche), forward and reverse Frame Shift Molecular Tags (FSMT) primers (Supplementary Table 2), and other necessary components, culminating in a total reaction volume of 50 µL. The FSMT primers, comprising six frames for both 338F and 806R types, were carefully mixed. To streamline the handling of multiple samples, a ‘ready-mix’ comprising essential Kapa components was prepared. This strategy was instrumental in reducing batch effects, particularly when utilizing reagents from different Kapa kits. The ready-mix was then allocated into aliquots each adequate for a 96-well plate. The first PCR cycle’s conditions were meticulously programmed to include specific durations and steps for denaturation, annealing, and extension, set at 95°C for one minute, followed by 10 cycles of 95°C for 15 seconds, 50°C for 30 seconds, and 72°C for 30 seconds, then 72°C for one minute, and a final indefinite hold at 4°C. Post-PCR, bead purification was performed to selectively exclude DNA fragments shorter than 300 bp, employing a 0.68:1 bead-to-DNA ratio. The second PCR phase (indexing) followed a similar methodology but with modifications in the reaction components, such as the Kapa HiFi Readymix (KAPA HiFi-Roche) and specific Adaptor_primer1 and indexing primers. The cycling conditions for this phase were set at 95°C for one minute, followed by 22 cycles of 95°C for 15 seconds, 60°C for 30 seconds, and 72°C for 30 seconds, concluding with 75°C for one minute and an indefinite hold at 4°C. Finally, for 16S sequencing, we used a series of custom-designed primers, chosen for their specificity and amplification efficiency. The sequences of these primers, along with the custom sequencing primer (NextF_Read1_seq), are comprehensively detailed in the supplementary material. Prepared libraries were pooled and sequenced on MiSeq Pair End 2×300 v2 flowcell (Supplementary Table 2).

#### 16S data processing and analysis

16S amplicon sequencing was performed using the moving pictures documentation in QIIME2.^17^ Briefly, sequences were imported into QIIME2 (version 2023.2). Paired end sequences were denoised with DADA2. Trimming values such as p-trim-left was set to 20 and p-trim-len was set to 220 and sampling depth was set at 120,000 sequences. Finally, an OTU table was produced for downstream analysis. Traditional microbial ecology analysis such as alpha diversity and beta diversity were also performed. In addition, taxonomic classification of the reads was performed in the QIIME2 using Silva 138 99% OTUs full-length sequences.

Relative abundances from operational taxonomic units (OTUs) were analyzed on the phylum, class, order, family, and genus levels. QIIME taxonomy labels with corresponding abundances were uploaded to MicrobiomeAnalyst, and the data were scaled by total sum prior to statistical and meta-analysis of microbiome data.^18^ Alpha diversity was accessed at the genus level by Chao, Shannon, and Simpson indexes with the Mann–Whitney test used to determine if alpha diversity differed across cohorts. Stacked bar graphs were used to visualize OTU relative abundances across cohorts. Pairwise comparisons between cohorts were made using the statistical method EdgeR. OTU correlations were analyzed using either Pearson’s r or Spearman’s rho correlation coefficients.

## Data Availability Statement

Data will be made available on NCBI via SRA

Metabolomics data files are accessible at MetaboLights (accession No. MTBLS10511).

## Acknowledgements

The authors would like to thank Katherine Havranek and Samia Dutra for stool DNA extractions. We gratefully acknowledge the technical support from the UNC Chapel Hill High Throughput Sequencing Facility (HTSF), a facility that is supported by the University Cancer Research Fund, Comprehensive Cancer Center Core Support grant (P30-CA016086).

## Author Contributions

KHM study coordination, recruitment of controls, sample preparation, DNA extractions, funding, and manuscript preparation. CJC collected and analyzed the final metabolomics data, provided bioinformatics for 16S data and cross-omics analysis, and conceptualized results. LR generated a list of microbial derived metabolites from the untargeted analysis, BAA performed stool metabolomics extractions, and ZAV processed the unidentified mass spectral features. PM next generation sequencing. BMW performed a pilot study with a subset of samples in this study. KAJ supervised the untargeted metabolomics analysis. CJC and SRC DHPS identification, characterization, pathway analysis, and conceptualized DHPS models in NDDs. The manuscript was written by CJC and KHM and reviewed by all authors. JCE, originator of MIND study and initial hypothesis, and conceptualization, study design, protocol development, 16S data analysis, manuscript preparation, and funding. RT, CMT, and RR-F screened, recruited, enrolled, and assessed patients and controls and edited the manuscript.

## Funding and Conflicts of Interest Statement

This study was sponsored by the Laboratory Directed Research and Development Program of Oak Ridge National Laboratory, managed by UT-Battelle, LLC, for the U.S. Department of Energy; the University of Tennessee’s College of Nursing and the Department of Chemistry; the University of Tennessee Office of Research, Innovation, and Economic Development’s Human

Health and Wellbeing Project; and by the Cole Family who support Parkinson’s care and research at the Cole Center for Parkinson’s Disease and Movement Disorders. The authors have no conflicts of interest to disclose.

Supplementary Information is available for this paper.

Correspondence and requests for materials should be addressed to Christopher Ellis.

Extended Data Figures are available.

