## Supplementary figures and images for "Specific Bacterial Taxa and Their Metabolite, DHPS, Linked to Alzheimer’s Disease, Parkinson’s Disease, and Amyotrophic Lateral Sclerosis"

### Abstract Art

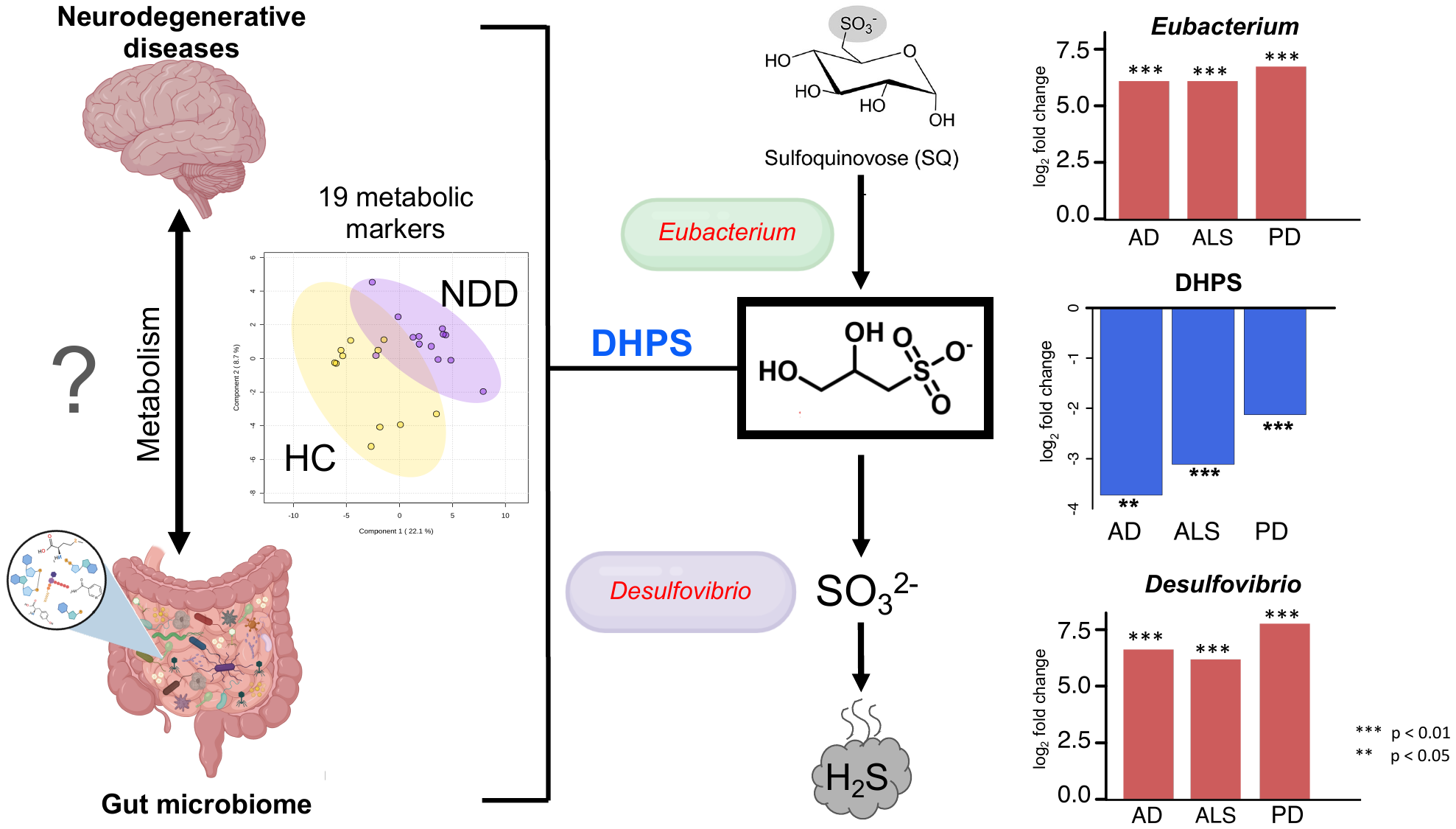
