## Extended Data for "Specific Bacterial Taxa and Their Metabolite, DHPS, Linked to Alzheimer’s Disease, Parkinson’s Disease, and Amyotrophic Lateral Sclerosis"

Extended Data Figure 1. Metabolites correlated with DHPS in NDDs.

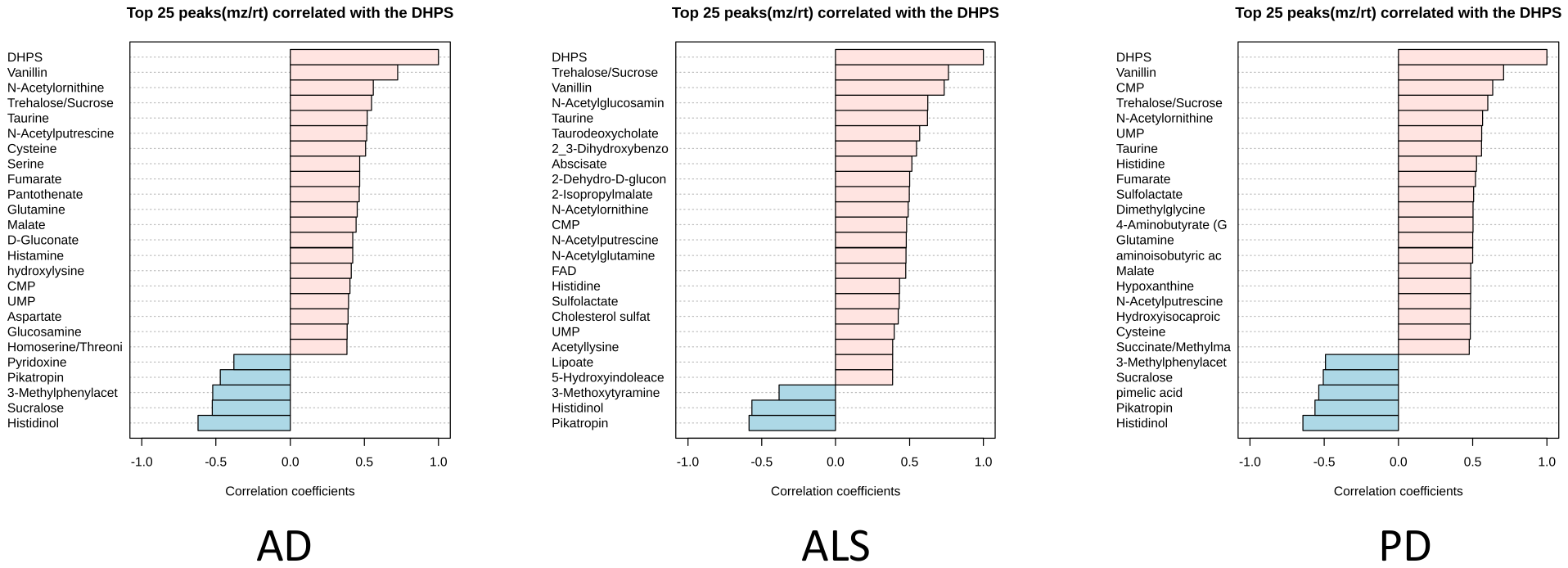

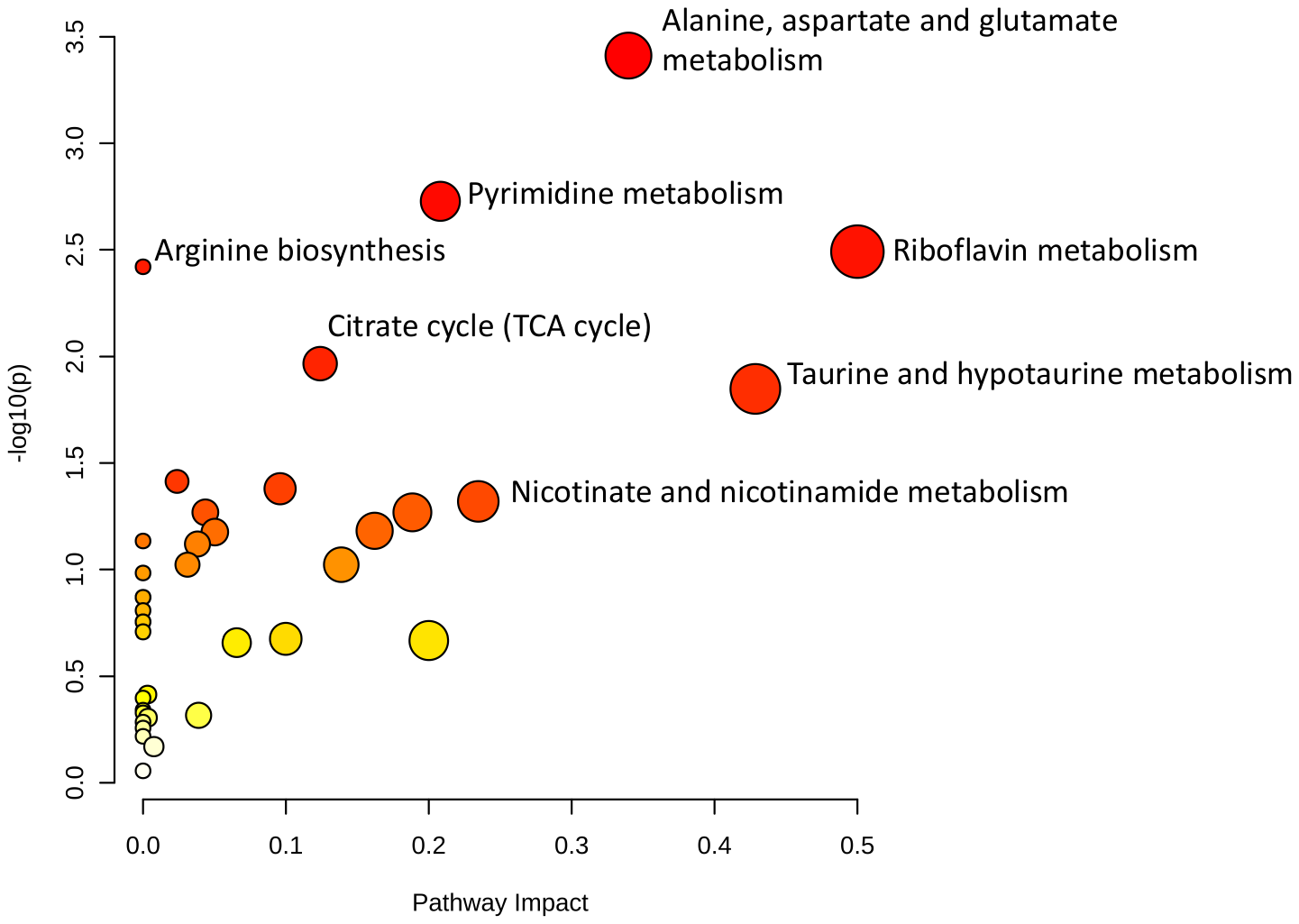

Extended Data Figure 2. Pathways altered in the AD stool metabolome using metabolites with VIP scores >1.

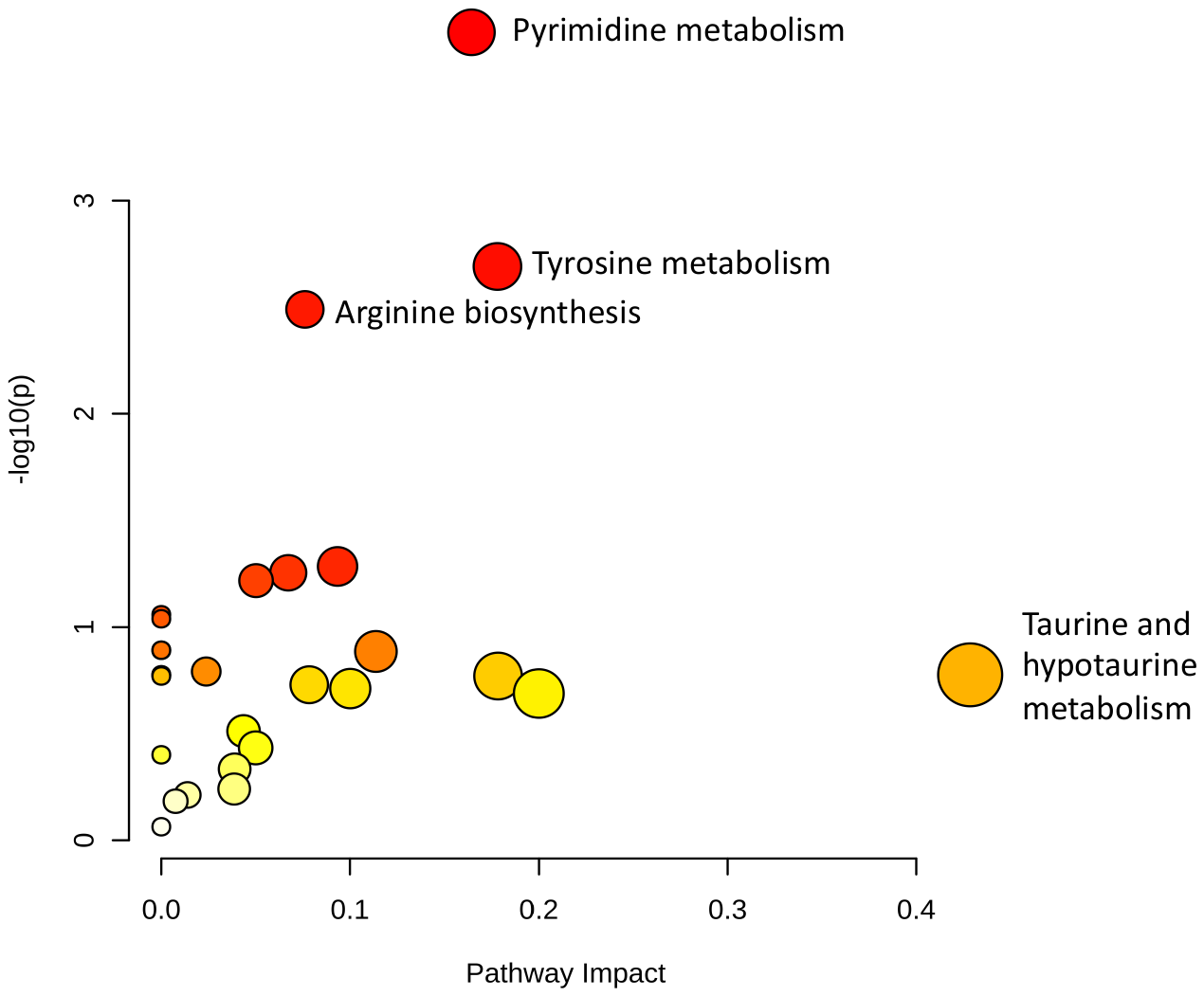

Extended Data Figure 3. Pathways altered in the ALS stool metabolome using metabolites with VIP scores >1.

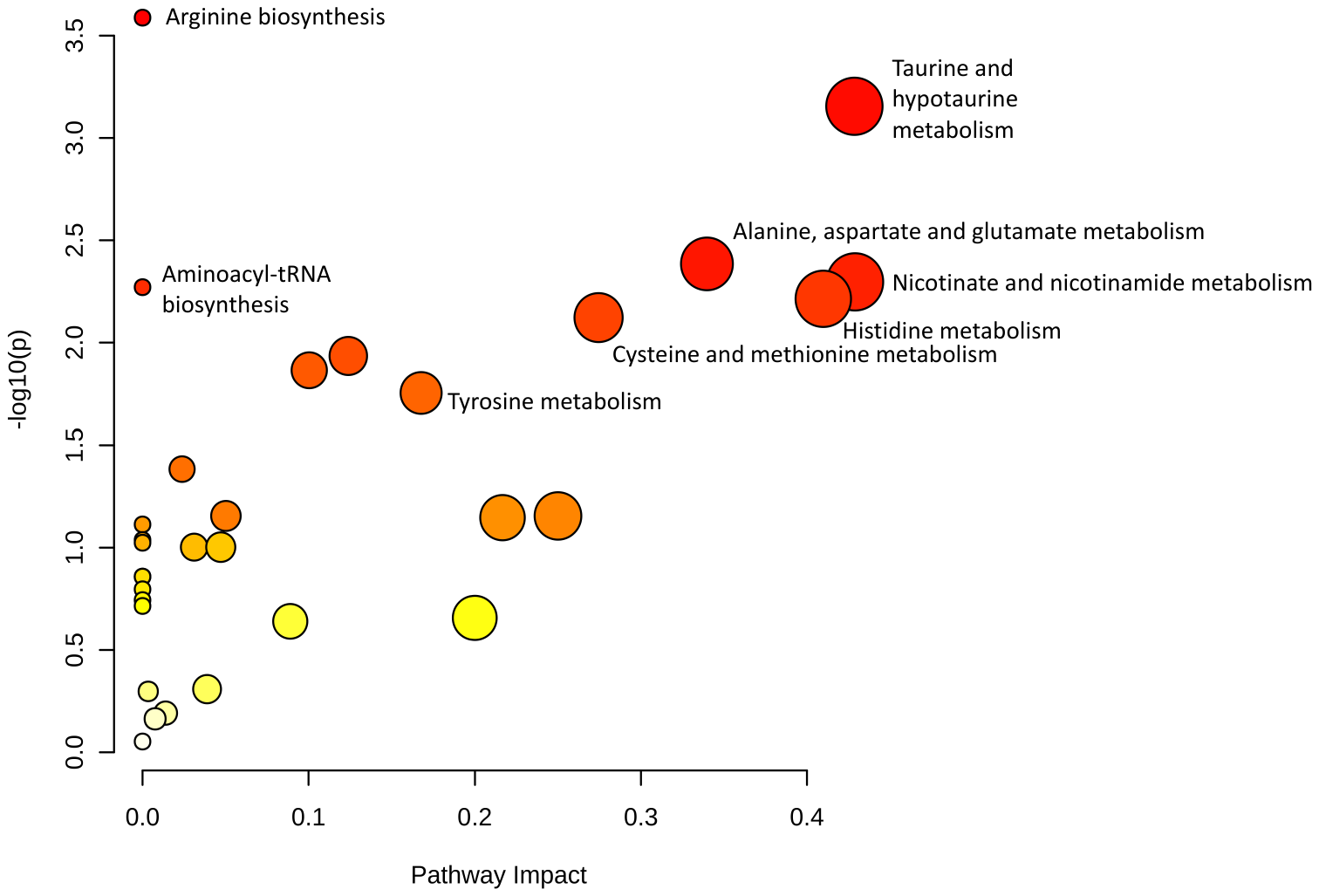

Extended Data Figure 4. Pathways altered in the PD stool metabolome using metabolites with VIP scores >1.

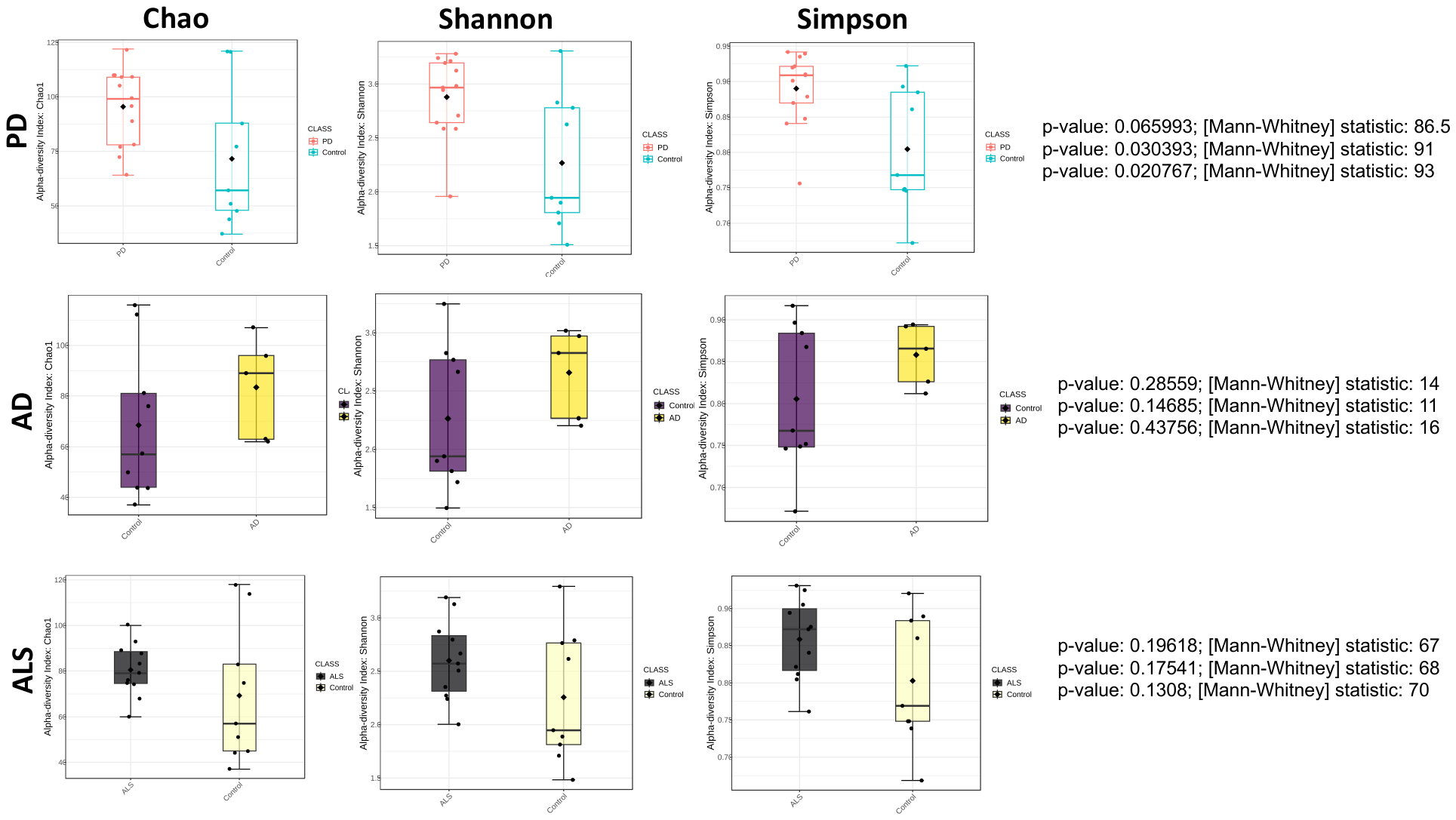

Extended Data Figure 5. Alpha diversity of stool microbiome in each NDD.

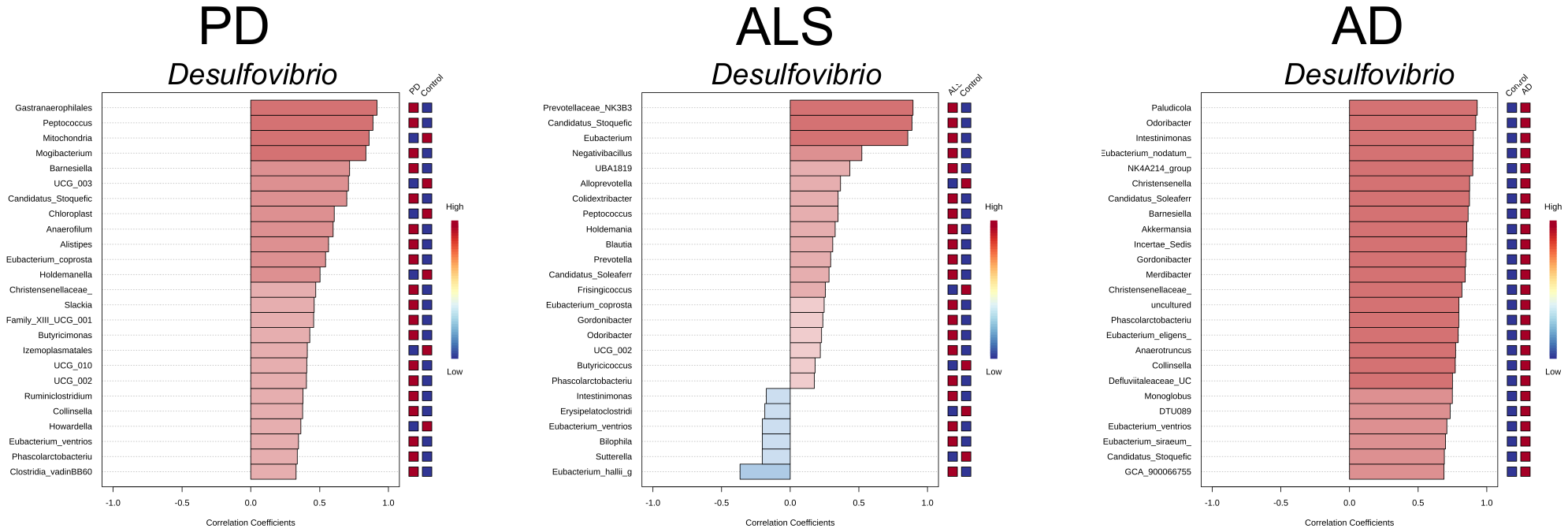

Extended Data Figure 6. Taxa correlated with *Desulfovibrio* in each NDD.

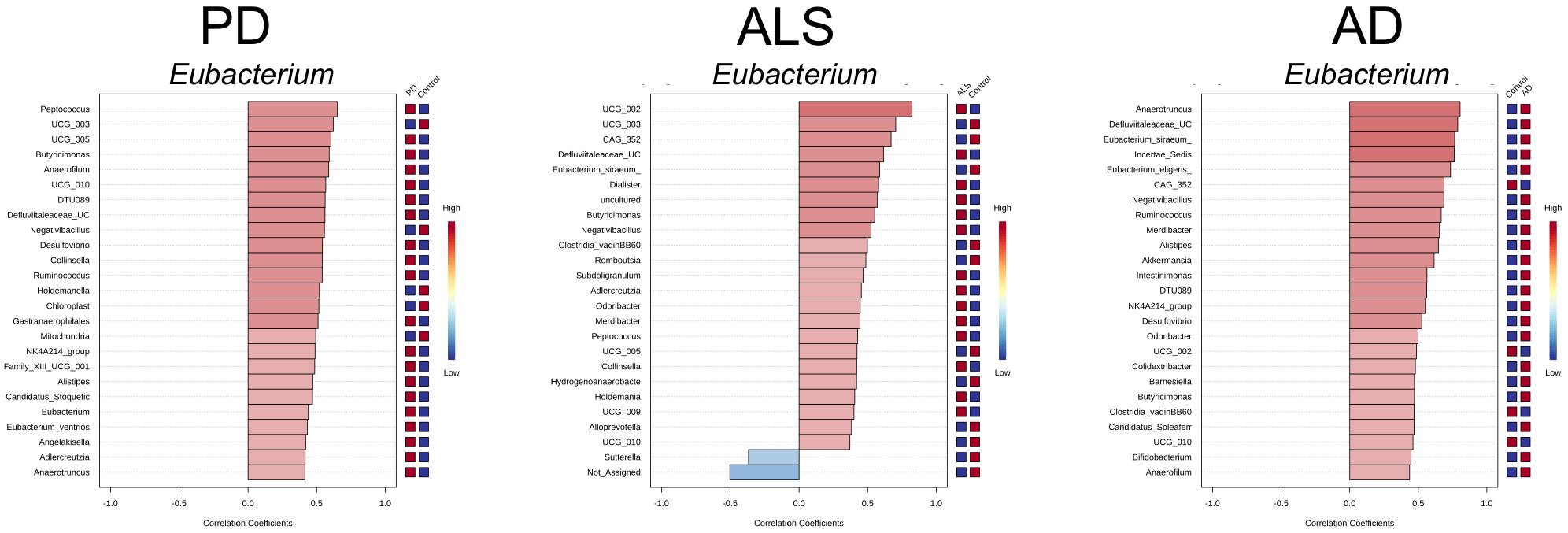

Extended Data Figure 7. Taxa correlated with *Eubacterium* in each NDD.

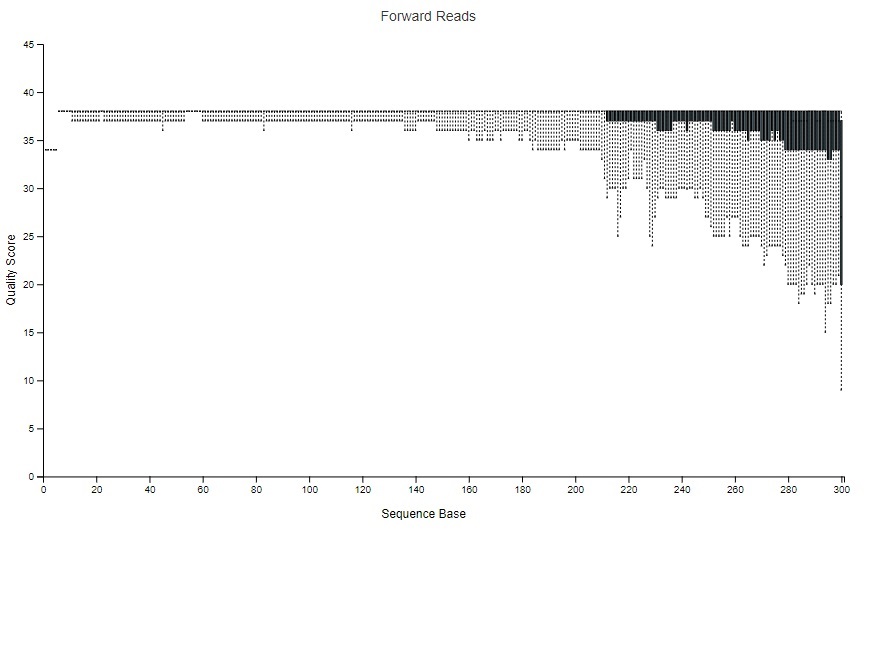

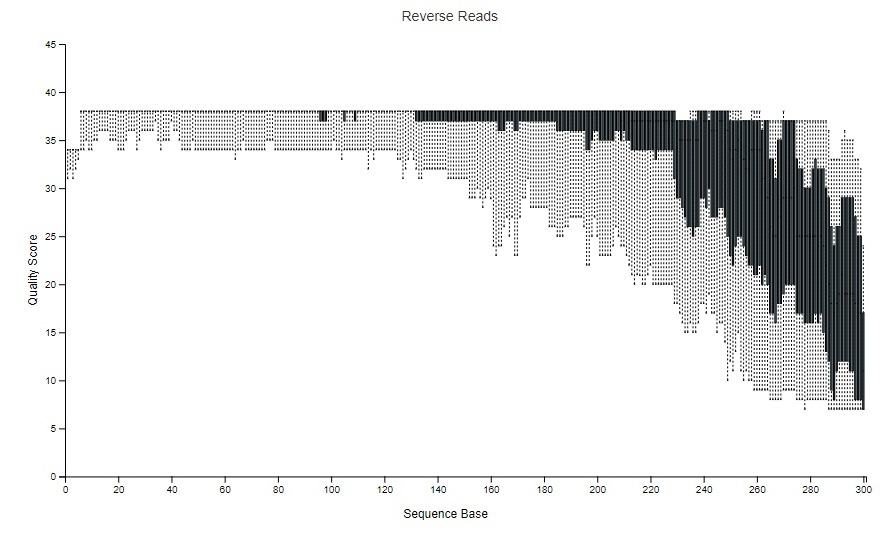

**Figure 8**: Amyotrophic lateral sclerosis (ALS) sequence quality plots prior to trimming. For ALS, p-trim-left was set to 20 and p-trim-len was set to 220.

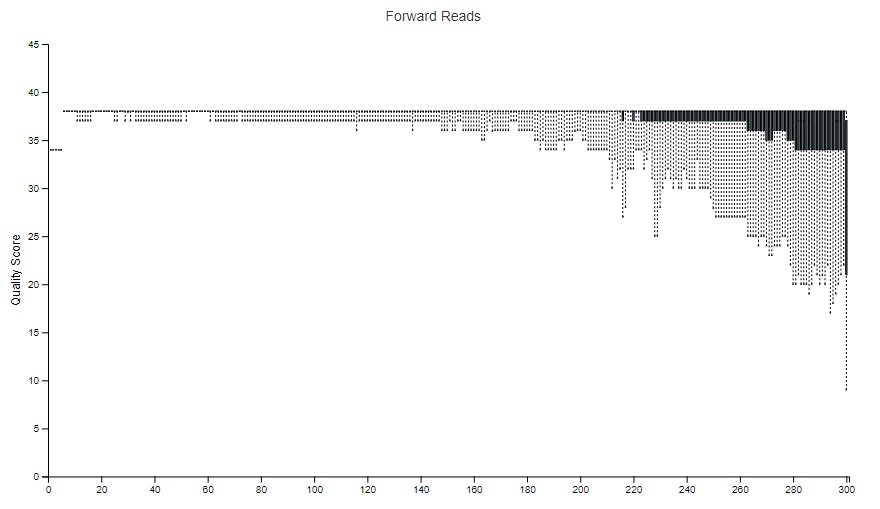

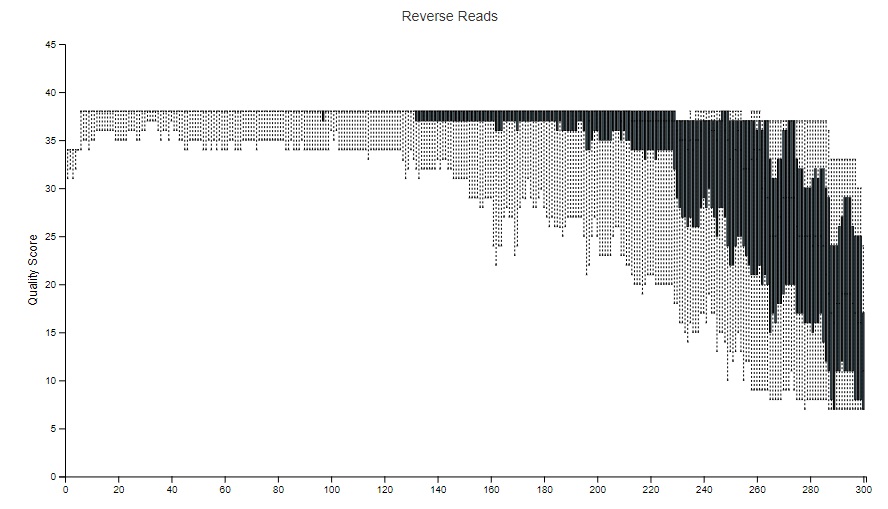

**Figure 9**: Alzheimer’s Disease (AD) sequence quality plots prior to trimming. For AD, p-trim-left was set to 20 and p-trim-len was set to 220.

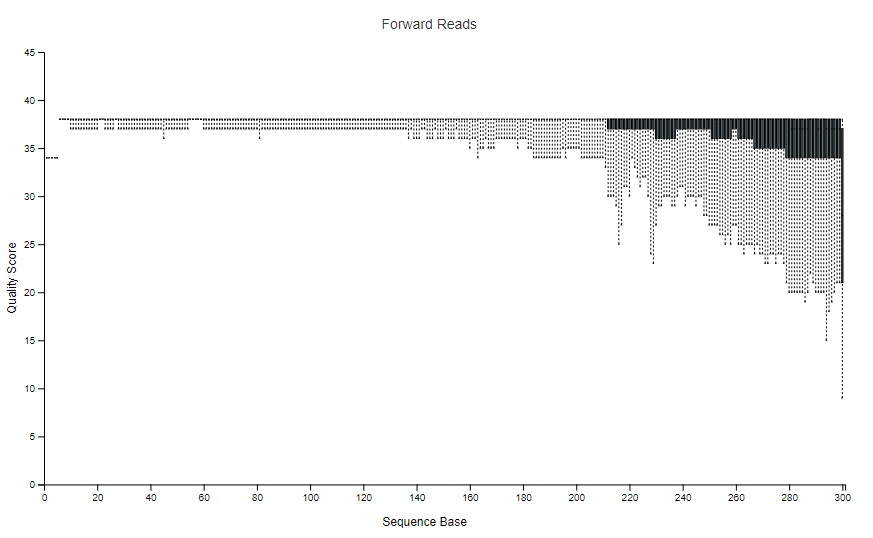

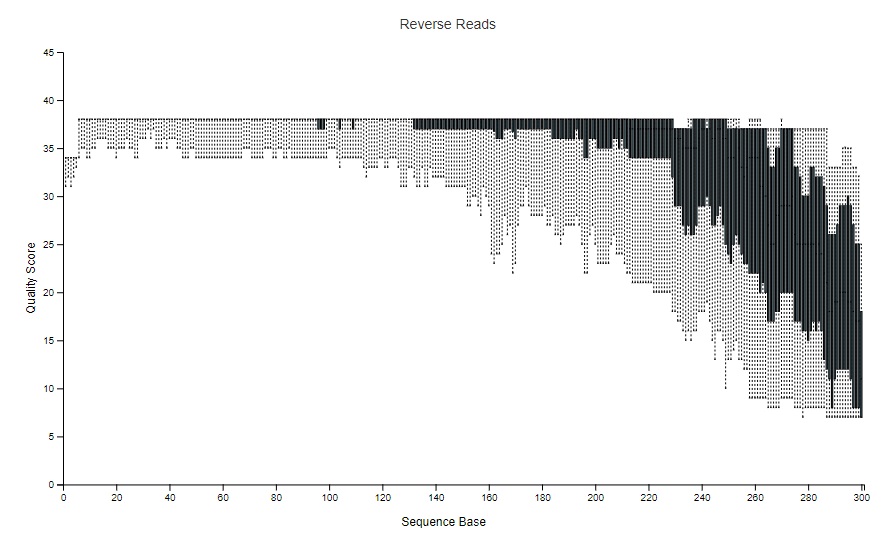

**Figure 10**: Parkinson’s Disease (PD) sequence quality plots prior to trimming. For PD, p-trim-left was set to 20 and p-trim-len was set to 220.
