## Supplementary for "Specific Bacterial Taxa and Their Metabolite, DHPS, Linked to Alzheimer’s Disease, Parkinson’s Disease, and Amyotrophic Lateral Sclerosis"

**SUPPLEMENTARY INFORMATION**

**Tables and Discussion**

**Supplementary Tables**

Supplementary Table 1. Demographics of Study Participants

|  | ALS | AD | PD | HC |
| --- | --- | --- | --- | --- |
| Number | 12 | 5 | 13 | 14 |
| Age in years mean (range) | 72.5 (71-74) | 64.5 (43-77) | 65.5 (51-80) | 66.1 (45-87) |
| Female | 4 | 3 | 9 | 8 |
| Male | 8 | 2 | 4 | 6 |
| BMI mean (range) | 25.4 (17.7-28.8) | 27.7 (19.2-37.8) | 28.3 (22.4-39.5) | 30.1(20-58.4) |
| Taking medications for NDD | 69% | 100% | 7% | 0 |
| Months from symptom onset (range) | 27.2 (6-82) | 35.7 (30-117) | 31.6 (15-57) |  |
| Months from diagnosis (range) |  | 18.3 (9-68) | 3 (0-20) |  |
| MOCA score |  | 19.8 |  | 26.5 |
| Clinical Dementia Rating Scale raw |  | 2.75 |  | 0 |
| Clinical Dementia Rating Scale global |  | 0.42 |  | 0 |
| ALS bulbar | 23% |  |  |  |
| ALS spinal | 77% |  |  |  |
| UPDRS-part I |  |  | 6.9 |  |
| UPDRS-part II |  |  | 6.7 |  |
| UPDRS-part III |  |  | 33.9 |  |
| UPDRS-part IV |  |  | 0 |  |
| UPDRS gastrointestinal symptoms |  |  | 0.23 |  |
| Hoehn and Yahr |  |  | 2.0 |  |

AD=Alzheimer’s Disease, ALS=amyotrophic lateral sclerosis, and PD=Parkinson’s Disease

**Supplementary Table 2. 16s Primers and Sequences**

| Primer | Sequence |
| --- | --- |
| 338F_f1_bc1 | TCCCTCGCGCCATCAGAGATGTGTATAAGAGACAGNNNNTGANNNNTCACTCCTACGGGAGGCAGCA |
| 338F_f2_bc1 | CCCTCGCGCCATCAGAGATGTGTATAAGAGACAGNNNNTTGANNNNTCACTCCTACGGGAGGCAGCA |
| 338F_f3_bc1 | TCCCTCGCGCCATCAGAGATGTGTATAAGAGACAGNNNNCTTGANNNNTCACTCCTACGGGAGGCAGCA |
| 338F_f4_bc1 | TCCCTCGCGCCATCAGAGATGTGTATAAGAGACAGNNNNACTTGANNNNTCACTCCTACGGGAGGCAGCA |
| 338F_f5_bc1 | TCCCTCGCGCCATCAGAGATGTGTATAAGAGACAGNNNNGACTTGANNNNTCACTCCTACGGGAGGCAGCA |
| 338F_f6_bc1 | TCCCTCGCGCCATCAGAGATGTGTATAAGAGACAGNNNNTGACTTGANNNNTCACTCCTACGGGAGGCAGCA |
| JMPM_806R_C | GTGACTGGAGTTCAGACGTGTGCTCTTCCGATCTNNNNCTAGGACTACHVGGGTWTCTAAT |
| JMPM_806R_2T | GTGACTGGAGTTCAGACGTGTGCTCTTCCGATCTNNNNTCTAGGACTACHVGGGTWTCTAAT |
| JMPM_806R_2A | GTGACTGGAGTTCAGACGTGTGCTCTTCCGATCTNNNNATCTAGGACTACHVGGGTWTCTAAT |
| JMPM_806R_G | GTGACTGGAGTTCAGACGTGTGCTCTTCCGATCTNNNNGGACTACHVGGGTWTCTAAT |
| JMPM_806R_A | GTGACTGGAGTTCAGACGTGTGCTCTTCCGATCTNNNNAGGACTACHVGGGTWTCTAAT |
| JMPM_806R_T | GTGACTGGAGTTCAGACGTGTGCTCTTCCGATCTNNNNTAGGACTACHVGGGTWTCTAAT |
| Adaptor_prim1 | AATGATACGGCGACCACCGAGATCTACACGCCTCCCTCGCGCCATCAGAGATGTG |
| PCR Primer, Index X | CAAGCAGAAGACGGCATACGAGAT**XXXXXX**GTGACTGGAGTTCAGACGTGTGCTC |
| NextF_Read1_seq | GCCTCCCTCGCGCCATCAGAGATGTGTATAAGAGACAG |

Supplementary Table 3: Denoised Dada2 sequence statistics for ALS.

| **sample-id** | **input** | **filtered** | **percentage of input passed filter** | **denoised** | **non-chimeric** | **percentage of input non-chimeric** |
| --- | --- | --- | --- | --- | --- | --- |
| UT001 | 3333341 | 3223084 | 96.69 | 3215930 | 2675917 | 80.28 |
| UT002 | 264625 | 248521 | 93.91 | 247619 | 230705 | 87.18 |
| UT004 | 146448 | 130887 | 89.37 | 130530 | 123611 | 84.41 |
| UT005 | 289498 | 272664 | 94.19 | 271792 | 246491 | 85.14 |
| UT007 | 332208 | 310075 | 93.34 | 309144 | 291650 | 87.79 |
| UT009 | 304345 | 285062 | 93.66 | 284213 | 267530 | 87.9 |
| UT021 | 283630 | 266267 | 93.88 | 265069 | 248373 | 87.57 |
| UT022 | 315324 | 296465 | 94.02 | 295263 | 280083 | 88.82 |
| UT024 | 281557 | 264174 | 93.83 | 263344 | 250738 | 89.05 |
| UT025 | 368934 | 345718 | 93.71 | 344637 | 316612 | 85.82 |
| UT026 | 371513 | 349092 | 93.96 | 347847 | 328903 | 88.53 |
| UT027 | 239266 | 145559 | 60.84 | 144093 | 139183 | 58.17 |
| UT029 | 271310 | 254847 | 93.93 | 253717 | 231167 | 85.2 |
| UT030 | 277108 | 261399 | 94.33 | 260379 | 231942 | 83.7 |
| UT031 | 339879 | 320361 | 94.26 | 319653 | 290788 | 85.56 |
| UT033 | 400676 | 375729 | 93.77 | 374342 | 348760 | 87.04 |
| UT035 | 304667 | 287292 | 94.3 | 286415 | 267373 | 87.76 |
| UT038 | 282330 | 263854 | 93.46 | 263209 | 247574 | 87.69 |
| UT043 | 337674 | 304102 | 90.06 | 302675 | 283800 | 84.05 |
| UT044 | 240100 | 218642 | 91.06 | 216363 | 209978 | 87.45 |
| UT045 | 379638 | 354653 | 93.42 | 352564 | 340359 | 89.65 |

Supplementary Table 4: Denoised Dada2 sequence statistics for AD.

| **sample-id** | **input** | **filtered** | **percentage of input passed filter** | **denoised** | **non-chimeric** | **percentage of input non-chimeric** |
| --- | --- | --- | --- | --- | --- | --- |
| UT001 | 3333341 | 3223084 | 96.69 | 3215930 | 2527333 | 75.82 |
| UT004 | 146448 | 130887 | 89.37 | 130530 | 124618 | 85.09 |
| UT005 | 289498 | 272664 | 94.19 | 271792 | 244145 | 84.33 |
| UT024 | 281557 | 264174 | 93.83 | 263344 | 244656 | 86.89 |
| UT031 | 339879 | 320361 | 94.26 | 319653 | 290788 | 85.56 |
| UT033 | 400676 | 375729 | 93.77 | 374342 | 339801 | 84.81 |
| UT036 | 319972 | 300338 | 93.86 | 299350 | 275837 | 86.21 |
| UT038 | 282330 | 263854 | 93.46 | 263209 | 244238 | 86.51 |
| UT039 | 320994 | 301967 | 94.07 | 301137 | 285524 | 88.95 |
| UT040 | 331360 | 311782 | 94.09 | 310839 | 280192 | 84.56 |
| UT041 | 277539 | 261496 | 94.22 | 260403 | 236380 | 85.17 |
| UT042 | 292256 | 275274 | 94.19 | 274322 | 257175 | 88 |
| UT043 | 337674 | 304102 | 90.06 | 302675 | 278357 | 82.43 |
| UT044 | 240100 | 218642 | 91.06 | 216363 | 204172 | 85.04 |
| UT045 | 379638 | 354653 | 93.42 | 352564 | 341915 | 90.06 |

Supplementary Table 5: Denoised Dada2 sequence statistics for PD.

| **sample-id** | **input** | **filtered** | **percentage of input passed filter** | **denoised** | **non-chimeric** | **percentage of input non-chimeric** |
| --- | --- | --- | --- | --- | --- | --- |
| UT001 | 3333341 | 3223084 | 96.69 | 3215930 | 2689852 | 80.7 |
| UT003A | 327352 | 308036 | 94.1 | 306966 | 295930 | 90.4 |
| UT004 | 146448 | 130887 | 89.37 | 130530 | 126167 | 86.15 |
| UT005 | 289498 | 272664 | 94.19 | 271792 | 246133 | 85.02 |
| UT011 | 402573 | 378535 | 94.03 | 377640 | 360867 | 89.64 |
| UT012 | 331555 | 311317 | 93.9 | 310091 | 286317 | 86.36 |
| UT013 | 386888 | 360794 | 93.26 | 359608 | 338338 | 87.45 |
| UT014 | 349287 | 327731 | 93.83 | 326512 | 306906 | 87.87 |
| UT015 | 371556 | 350038 | 94.21 | 348850 | 339371 | 91.34 |
| UT016 | 348217 | 328968 | 94.47 | 328013 | 303594 | 87.19 |
| UT018 | 342045 | 321908 | 94.11 | 320906 | 298046 | 87.14 |
| UT019 | 305939 | 287747 | 94.05 | 286866 | 279062 | 91.21 |
| UT023 | 378100 | 355478 | 94.02 | 354324 | 340951 | 90.17 |
| UT024 | 281557 | 264174 | 93.83 | 263344 | 248938 | 88.41 |
| UT031 | 339879 | 320361 | 94.26 | 319653 | 291581 | 85.79 |
| UT033 | 400676 | 375729 | 93.77 | 374342 | 349541 | 87.24 |
| UT038 | 282330 | 263854 | 93.46 | 263209 | 247674 | 87.73 |
| UT043 | 337674 | 304102 | 90.06 | 302675 | 286359 | 84.8 |
| UT044 | 240100 | 218642 | 91.06 | 216363 | 203626 | 84.81 |
| UT045 | 379638 | 354653 | 93.42 | 352564 | 340822 | 89.78 |
| UT060 | 518396 | 468026 | 90.28 | 466205 | 455960 | 87.96 |
| UT061 | 397674 | 372754 | 93.73 | 371220 | 361173 | 90.82 |
| UT063 | 348004 | 323784 | 93.04 | 322503 | 314395 | 90.34 |

Supplementary Table 6. Comprehensive List of Metabolites with FC>1.5 and p<0.1 and/or VIP>1

| **AD** | **ALS** | **PD** |
| --- | --- | --- |
| DHPS | N-Acetylglucosamine 1/6-phosphate | Trehalose/Sucrose |
| Fumarate | Cholesterol sulfate | 2-Isopropylmalate |
| Malate | 2-Isopropylmalate | Phenyllactic acid |
| 4-Pyridoxate | Xanthurenic acid | 5-Hydroxyindoleacetic acid (5-HIAA) |
| Lipoate | Kynurenic acid | Sulfolactate |
| UMP | Vanillin | Histidine |
| Biotin | DHPS | 2-Hydroxy-2-methylsuccinate |
| IMP | Glycodeoxycholate | Lipoate |
| Deoxyribose phosphate | N-Acetylglutamine | 2-hydroxyglutaric acid |
| Cysteine | Sulfolactate | 2-Dehydro-D-gluconate |
| D-Gluconate | 2_3-Dihydroxybenzoate | UMP |
| Aspartate | Lipoate | CMP |
| Taurine | 3_4-Dihydroxyphenylacetate (DOPAC) | DHPS |
| FAD | 5-Hydroxyindoleacetic acid (5-HIAA) | D-Gluconate |
| Histamine | D-Glucarate | N-Acetylglucosamine 1/6-phosphate |
| hydroxylysine | N-Acetylornithine | Biotin |
| Glucosamine phosphate | Pyridoxine | Cytidine |
| S-Ribosyl-L-homocysteine | Aconitate | 3-Methylthiopropionate |
| CMP | 2-Dehydro-D-gluconate | Fumarate |
| N-Acetylglucosamine 1/6-phosphate | Jasmonate | Hypoxanthine |
| homocarnosine | Taurodeoxycholate | Aspartate |
| Vanillin | FAD | Gluconolactone |
| Pikatropin | Sedoheptulose 1/7-phosphate | xylose |
| 3-Methylthiopropionate | Creatine | Cholesterol sulfate |
| Hypoxanthine | sn-Glycerol 3-phosphate | N-Acetylornithine |
| Aconitate | UMP | Glutamine |
| Histidinol | Dopamine | N-Acetylglucosamine |
| Serine | 2-Hydroxy-2-methylsuccinate | Vanillin |
| D-Glyceraldehdye 3-phosphate | Creatinine | Dopamine |
| Glycerone phosphate | Homovanillic acid (HVA) | glutaric acid |
| dTMP | 2-hydroxyglutaric acid | methyl succinic acid |
| Homovanillic acid (HVA) | CMP | Homovanillic acid (HVA) |
| Cholesterol sulfate | Trehalose/Sucrose | N-Acetyl-beta-alanine |
| 2-Dehydro-D-gluconate | methyl succinic acid | Malate |
| 3-Methylphenylacetic acid | glutaric acid | Aconitate |
| Glutamine | Glutamine | Histidinol |
| Guanine | N-Acetylputrescine | Taurine |
| N-Carbamoyl-L-aspartate | Asparagine | Cystathionine |
| Trehalose/Sucrose | Orotate | 3_4-Dihydroxyphenylacetate (DOPAC) |
| sn-Glycerol 3-phosphate | Cytidine | Lysine |
| D-Glucarate | N-Acetylglucosamine | NAD+ |
| Sulfolactate | Histidinol | Xylitol |
| Carnitine | Taurine | Serine |
| Phenyllactic acid | CDP-choline | Sucralose |
| FMN | Guanosine | IMP |
| Sucralose | Cystathionine | Histamine |
| NAD+ | Biotin | Taurodeoxycholate |
|  | Pikatropin | GMP |
|  | Arginine | Hydroxyisocaproic acid |
|  | Uridine | 3-Hydroxyisovalerate |
|  | GMP | pimelic acid |
|  | 3-Methoxytyramine (3-MT) | Cysteate |
|  | Hydroxyisocaproic acid | Pikatropin |
|  | 3-Methylthiopropionate | Cysteine |
|  | Ribose phosphate | Nicotinamide |
|  | Abscisate | Asparagine |
|  |  | FAD |

AD=Alzheimer’s Disease, ALS=amyotrophic lateral sclerosis, and PD=Parkinson’s Disease

SupplementaryTable 7. Biomarkers (Metabolites) Unique to AD, ALS, and PD

| **Unique to ALS** | **Unique to AD** | **Unique to PD** | **Conserved in**  **ALS, AD, PD** | **ALS and PD** | **ALS and AD** | **AD and PD** |
| --- | --- | --- | --- | --- | --- | --- |
| Kynurenic acid | hydroxylysine | Histidine | N-Acetylglucosamine 1/6-phosphate | 2-Isopropylmalate | D-Glucarate | IMP |
| Xanthurenic acid | 4-Pyridoxate | Gluconolactone | Cholesterol sulfate | Taurodeoxycholate | sn-Glycerol 3-phosphate | Histamine |
| 2_3-Dihydroxybenzoate | Deoxyribose phosphate | xylose | DHPS | 5-Hydroxyindoleacetic acid (5-HIAA) |  | Cysteine |
| Glycodeoxycholate | Glucosamine phosphate | 3-Hydroxyisovalerate | CMP | Cystathionine |  | Malate |
| Sedoheptulose 1/7-phosphate | S-Ribosyl-L-homocysteine | pimelic acid | Vanillin | N-Acetylornithine |  | Fumarate |
| Creatine | homocarnosine | N-Acetylglucosamine | Aconitate | Asparagine |  | D-Gluconate |
| Creatinine | Serine | Cysteate | UMP | Cytidine |  | Hypoxanthine |
| CDP-choline | D-Glyceraldehdye 3-phosphate | Xylitol | Sulfolactate | Dopamine |  | Aspartate |
| N-Acetylglutamine | Glycerone phosphate | Nicotinamide | Histidinol | 2-Hydroxy-2-methylsuccinate |  | Phenyllactic acid |
| Guanosine | dTMP | Lysine | Taurine | GMP |  | Sucralose |
| DOPAC | 3-Methylphenylacetic acid |  | Homovanillic acid (HVA) | 2-hydroxyglutaric acid |  |  |
| Orotate | Guanine |  | FAD | Hydroxyisocaproic acid |  |  |
| N-Acetylputrescine | N-Carbamoyl-L-aspartate |  | 2-Dehydro-D-gluconate | methyl succinic acid |  |  |
| Jasmonate | Carnitine |  | Biotin | glutaric acid |  |  |
| Arginine | FMN |  | Lipoate |  |  |  |
| Uridine |  |  | Pikatropin |  |  |  |
| 3-Methoxytyramine (3-MT) |  |  | Glutamine |  |  |  |
| Pyridoxine |  |  | 3-Methylthiopropionate |  |  |  |
| Ribose phosphate |  |  | Trehalose/Sucrose |  |  |  |
| Abscisate |  |  |  |  |  |  |

[AD=Alzheimer’s Disease, ALS=amyotrophic lateral sclerosis, and PD=Parkinson’s Disease]

**Supplementary Discussion**

***Melainabacteria could be associated with DHPS mediated cryptic sulfur metabolism in NDDs***

In 2013, a novel class of microbes, Melainabacteria, were discovered in the human gut.^26^ Melainabacteria are a non-photosynthetic clade of cyanobacteria and their role in the human gut and physiology is largely unknown.^26-29^ This is of particular interest in our study as SQDG is highly abundant in cyanobacteria. The abundance of Melainabacteria in gut is correlated with diet as the highest abundances of Melainabacteria in stool have been found in herbivore populations, with lower abundances in omnivore and carnivore populations suggesting that Melainabacteria have a role in fermenting dietary plant fibers.^26^ This may also provide another connection of DHPS to diet.

Furthermore, a partial metabolic reconstruction from Soo *et al*. (2014) of *Gastranaerophilales,* one of the six major taxonomic orders within the Melainabacteria class, is predicted to use the Embden–Meyerhof pathway to acquire energy by fermenting simple carbohydrates, and DHPS was positively correlated with trehalose in this study suggesting a link between DHPS and *Gastranaerophilales*.^28^ This metabolic reconstruction also revealed that *Gastranaerophilales* have the genes required for the biosynthesis of B vitamins (biotin, riboflavin, nicotinamide, and dihydrofolate) and vitamin K which could demonstrate a beneficial relationship between Melainabacteria and humans.^26,28,30^ The basis for the symbiotic relationship between *Thalassiosira pseudonana* and Roseobacters in aquatic systems was vitamin B_12_ and DHPS.^31^ Given that Melainabacteria are suggested to have a syntrophic H_2_-producing niche, it is likely these microbes can degrade dietary sulfonates. In addition, a previous study has shown that *Gastranaerophilales* is one of the main producers of microbial-derived indole leading to increased indole in PD.^32^ We found that DHPS was negatively correlated with both biotin and metabolites involved in indole metabolism. The majority of preliminary studies show a trend of increased Melainabacteria abundance in patients with metabolic, neurodegenerative, and gastrointestinal disease, though results in literature are still inconsistent as understanding the role of *Gastranaerophilales* in the human gut remains in its infancy.^30,33^ In this study, we found that the abundance of *Gastranaerophilales* was increased in PD, ALS, and AD compared to the HC cohort. This contradictory evidence highlights the need to further investigate the role of Melainabacteria. In addition, *Gastranaerophilales* was strongly correlated to *Desulfovibrio.* This may suggest that *Gastranaerophilales* plays a role in DHPS metabolism since *Desulfovibrio* has the cellular machinery to both produce and degrade DHPS. With this, we propose there could be a relationship between Melainabacteria and DHPS given the phylogenetic similarity to cyanobacteria and the negative correlation between DHPS and biotin.

**Metabolome alterations in NDDs**

While it has become widely accepted that gut dysbiosis is a contributing factor in NDDs, the pathophysiological mechanism remains elusive. This is, in part, due to the complexity of the gut microbiome and metabolome and “dark matter” in host-microbiome interactions. This “dark matter” represents the vast amount of detected, but unidentified, small molecules in the metabolome. These analytes likely impact human physiology, highlighting the need for better understanding the small molecules involved in host-microbiome interactions in the gut. To this end, we used an untargeted, hypothesis-generating approach to investigate gut microbiome alterations in NDDs.

As expected, we observed differences in the gut microbiome composition and function in patients with NDDs (AD, ALS, and PD) compared to HCs. The presented data align with previous findings in the literature that sulfur metabolism, tryptophan catabolism, bile acids, and amino acids are altered in NDDs. ^1-4^ We identified 19 consistent markers for AD, ALS, and PD, as well as metabolic signatures unique to each NDD.

Carnitine was a metabolic signature unique to AD and is known to play a crucial role in energy metabolism and mitochondrial function. Dietary-carnitine supplements have been proposed for prevention or alleviation of AD. ^5^ In ALS, creatine and creatinine were unique signatures and decreased in ALS. Creatine has been proposed to exhibit protective effects against mitochondrial dysfunction and may also be neuroprotective as it has been linked to increased survival time in an ALS clinical trial. ^6^

Two of the unique metabolic signatures for PD were decreased xylose and xylitol, indicating impaired sugar metabolism. Xylose and xylitol feed into the pentose phosphate pathway (PPP) resulting in the production of NADPH, pyruvate, and shikimate, which are building blocks for some branched chain and aromatic amino acids and antioxidants. This result is compelling since neurons preferentially use the PPP for sugar metabolism and rely on the metabolic products of the PPP for protection against oxidative stress. ^7^ Decreased flux through the PPP results in reduced antioxidant capacity and increased oxidative stress, which is a leading hypothesis for pathogenesis of PD. ^7^

The 19 metabolic markers for NDD showed significant alterations in sulfur, cholesterol, sugar, and vitamin B metabolism. To our surprise, one of these markers was DHPS, a sulfonated microbial derived metabolite known for its role in sulfur and carbon flux in marine ecosystems, with an unknown role in human physiology. ^8,9^ With this, we have discovered a novel link between microbial derived metabolite, DHPS, and NDDs as DHPS was a significant driver of global metabolome differences when comparing NDD and HC cohorts. To the best of our knowledge, we are the first to provide data linking DHPS to three NDDs: AD, ALS, and PD, thereby, demonstrating a potential role of DHPS in human physiology.
